# Engineering Functional CLA-Targeting CAR Approaches for Pancreatic Ductal Adenocarcinoma

**DOI:** 10.64898/2026.07.03.736395

**Authors:** Camille Dourlens, Kim Vanderliek, Lilli Geiger, Nico Burzan, Stefan Tomiuk, Miriam Droste, Andre Felsberger, Helene Hubrich, Jana Winkler, Olaf Hardt, Daniel Schäfer

## Abstract

Pancreatic cancer remains a highly lethal malignancy with limited therapeutic options. Chimeric antigen receptor (CAR) therapy has revolutionized the treatment of hematological cancers but still faces major limitations in solid tumors, particularly due to the scarcity of tumor-specific targets. Cutaneous lymphocyte antigen (CLA) recently emerged as a promising PDAC target due to its high tumor expression and limited presence in healthy tissues. However, previously reported CLA-directed CAR constructs lacked antitumor functionality. Here, we investigated multiple strategies to generate functional CLA-targeting CAR approaches. We first hypothesized that impaired activity resulted from fratricide caused by CLA expression on activated T cells. CLA knockout was successfully achieved through deletion of fucosyltransferase-7, but not by knockout of the major CLA carrier backbones CD162, CD44 or CD43, suggesting additional CLA carriers or compensatory regulation. As CLA knockout alone did not restore CAR-mediated killing, we explored whether insufficient binding affinity limited CAR activity. Affinity maturation was performed *in silico* and *in vitro* using yeast surface display, identifying 39 candidate mutations, although none restored cytotoxicity. We finally switched to an AdCAR strategy using anti-biotin CAR T cells combined with biotinylated anti-CLA scFv-Fc adapters. This approach enabled efficient, concentration-dependent cytotoxicity with both CLA-targeting binders. Additionally, we identified a dynamic, cell density-dependent regulation of CLA expression. Finally, glycan profiling of CLA binders further revealed broader-than-expected glycan interactions, suggesting a potentially wider definition of the CLA family. Overall, our findings establish CLA as a functional PDAC immunotherapy target while revealing unexpected complexity in its regulation and molecular presentation.

## 1 Introduction

Pancreatic ductal adenocarcinoma (PDAC) is a highly lethal disease, currently the third leading cause of cancer-related death in the U.S. and Europe, accounting for ~8% of deaths despite only ~3% of cancer cases^1,2^. Therapeutic options remain limited to chemoradiotherapy and surgery, yet ~80% of patients present with unresectable disease, resulting in a poor ~13% 5-year survival rate^3,4^. This underscores the urgent need for novel therapeutic strategies^5^. Chimeric antigen receptor (CAR) T cells have transformed adoptive immunotherapy, becoming standard-of-care for multiple hematological malignancies^6^. These genetically engineered cells specifically recognize and eliminate target cells^7^. However, their efficacy in solid tumors remains limited, largely due to the challenge of identifying tumor-restricted antigens^8^. This highlights the importance of target selection for safe and effective CAR T cell therapy in solid tumors.

Recently, four promising PDAC-associated targets were identified: CD318, TSPAN8, CD66c and cutaneous lymphocyte antigen (CLA)^9^. CARs targeting each of these four targets have been generated, with CD318 advancing to a phase I/IIa clinical trial (NCT07153289). CLA is particularly attractive due to its favorable safety profile and restricted expression on a subset of peripheral leukocytes. Beyond PDAC, CLA is also overexpressed in lymphoma^10^. However, CLA-directed CARs have not yet demonstrated functional activity, warranting further investigation^9^.

CLA was described as a carbohydrate epitope expressed on circulating leukocytes that mediates skin homing^11^ and is defined by binding of the HECA-452 IgM antibody^12^. Although primarily associated with memory T cells, CLA is also expressed on memory B cells, neutrophils and innate lymphoid cells^13^. Depletion of specific immune subsets can be clinically managed, as demonstrated by CD19-targeting CAR T cell therapies^14^, making CLA still a promising target candidate. CLA biosynthesis depends on glycosylation mediated by glycosyltransferases, particularly fucosyltransferase VII (FUT7)^15^. Initially described on P-selectin glycoprotein ligand-1 (CD162/PSGL1/SELPLG)^11^, CLA has also been identified on CD44^16^ and CD43^17^. Structurally, CLA consists of sialyl Lewis X-related motifs with additional fucosylation and, in some reports, sulfation^18,19^, although its precise molecular definition remains incomplete. Functionally, CLA binds E- and P-selectin on dermal endothelial cells, enabling leukocyte tethering and rolling during extravasation, thereby promoting recruitment to inflamed skin^20^. Elevated CLA^+^ T cells frequencies are observed in psoriasis, atopic dermatitis and cutaneous T cell lymphoma^21^. The combination of CLA overexpression in PDAC, the availability of a well-characterized binding antibody, and its restricted expression on a dispensable subset of leukocytes collectively make CLA an attractive target for CAR T cell therapy. However, important biological aspects of CLA remain poorly understood, potentially limiting the development of effective CLA-directed cellular therapies.

In this study, we aimed to address the limitations preventing the development of functional CLA-directed CAR T cells. We identified a density-dependent regulation of CLA expression and investigated the underlying mechanisms. In parallel, we explored the structure and cellular localization of CLA, which remain incompletely defined and inconsistently described in literature.

## 2 Methods

### Cell lines and culture conditions

AsPC1, BxPC3, HL60, SupT1, HEK293T, PanCa0203 and Capan2 cells (ATCC) were cultured in RPMI or DMEM high-glucose medium (Biowest) supplemented with 2 mM L-glutamine and 10% fetal bovine serum (FBS; Eximus). OCI-AML2 (DSMZ) cells were cultured in Alpha-MEM Eagle medium (PAN-Biotech) with 20% FBS. PanCa0201 primary PDAC cell line was generated from dissociated human PDAC biopsies using the Tumor Dissociation Kit, gentleMACS™ Octo Dissociator with Heaters, and Tumor Cell Isolation Kit, followed by culture in Pancreas TumorMACS™ medium (Miltenyi Biotec). *S. cerevisiae* EBY100 (ATCC) were cultured in YPD, SDCAA or SGCAA media, as previously described^22^. For cell-density experiments, cells were seeded at defined densities and cultured under indicated conditions, including variable FBS concentrations, stress conditions, or supplementation with IL-12, IL-4 or TGF-β (Miltenyi Biotec). Marker expression was daily analyzed by flow cytometry.

### Flow cytometry

Samples were acquired on MACSQuant® Analyzer 10 or 16 instruments (Miltenyi Biotec) and analyzed using MACSQuantify™ (Miltenyi Biotec, v2.13.0), FlowJo (BD Biosciences, v10.10.0), GraphPad Prism (GraphPad Software, v10.1.2) and Microsoft Excel (Microsoft, v2502). Staining with Miltenyi Biotec antibody conjugates was performed according to manufacturer instructions. Dead cells were excluded using 7-AAD. All antibodies used in this study are listed in supplementary table 1. Gating strategies are shown in supplementary figure S1.

### Cyclic immunofluorescence

Primary human PDAC tissues were collected at the University Medical Center Göttingen from patients undergoing surgical resection of tumor masses. Fresh-frozen healthy organ tissue samples were purchased from ProteoGenex and BioIVT. Tissues were prepared, fixed and stained as previously described^9^. Imaging was performed on the MACSima™ Imaging Platform (Miltenyi Biotec) according to the manufacturer’s instructions. For figure display, all HDR images were uploaded in ImageJ (v1.49) and the histogram was set to equal values for all images from 0 to 29980.

### CRISPR/Cas9 knock-out (KO) generation

CD162, CD44, CD43 and FUT7 KO were generated by CRISPR-Cas9 electroporation using the CliniMACS® Electroporator integrated into the CliniMACS® Prodigy system (Miltenyi Biotec). The three recommended predesigned sgRNAs (AA, AB and AC), Cas9 Nuclease V3 and electroporation enhancer were obtained from IDT and used according to manufacturer recommendations. The KO in T cells was performed three days after isolation.

### CAR/AdCAR construct generation and lentiviral production

Synthetic gBlocks (IDT) were cloned into Miltenyi Biotec backbones containing LacZ and kanamycin resistance cassettes. Bve1 restriction enzymes, T4 ligase and cloning reagents were provided by NEB, according to their standard protocol. Plasmids were transformed into 5-alpha competent *E-coli* (NEB) and purified using Zyppy™ Miniprep and ZymoPURE™ Maxiprep kits (Zymo research). AdCAR constructs were generated as previously described^23^. Lentiviral (LV) particles were produced by transient polyethylenimine-mediated transfection of HEK293T cells. Viral supernatants were collected after 48 h, concentrated by overnight centrifugation, resuspended in TexMACS™ medium (Miltenyi Biotec) and stored at −70°C. Viral titers were determined by SupT1 transduction.

### T cell isolation and CAR/AdCAR transduction

Peripheral blood mononuclear cells (PBMCs) were isolated by PanColl (PanBiotech) density-gradient centrifugation from whole blood. T cells were purified using the Pan T Cell Isolation Kit and activated in TexMACS™ medium containing 1:100 TransAct™, IL-7 and IL-15 (12.5 ng/mL each; all Miltenyi Biotec). Cells were transduced with lentivirus 24h after activation. Three days later, TransAct™ was removed and cells were cultured in IL-7/IL-15-supplemented medium. Transduction efficiency was assessed by flow cytometry.

### Anti-CLA scFv-Fc production

Anti-CLA scFv-Fc expression constructs were synthesized, cloned into the constitutive mammalian expression vector pcDNA™3.4 TOPO™ and prepared by GeneArt (ThermoFisher Scientific). Expi293F cells were expanded and transiently transfected according to the manufacturer’s instructions using the ExpiFectamine™ 293 Transfection Kit (ThermoFisher Scientific). Four days post-transfection, cultures were clarified by addition of 100 g/L Celite 545 filter aid (Sigma-Aldrich) followed by sterile filtration through a 0.22 μm PES membrane (Nalgene™ Rapid-Flow™, ThermoFisher Scientific). Cell-free supernatants were stored at −20°C until purification. ScFv-Fc proteins were purified using a HiTrap™ MabSelect column (Cytiva) according to the manufacturer’s protocol and buffer-exchanged into PBS (pH 7.4) using PD-10 Sephadex™ G-25 desalting columns (Cytiva). Protein purity was assessed by SDS-PAGE under reducing conditions. Samples were mixed with 5× SDS sample buffer containing DTT, heated at 95°C for 5 min, and separated on 4-20% Criterion™ TGX™ Precast Midi Protein Gels (Bio-Rad). Electrophoresis was performed at 220 V for 50 min, followed by Coomassie Blue staining for 30 min at room temperature.

### Adapter biotinylation

Pure antibodies (Miltenyi Biotec) and scFv-Fc molecules were buffer-exchanged into 0.1 M NaHCO3 solution (Sigma-Aldrich) via NAP-5 columns (Cytiva). Proteins were biotinylated for 1h at room temperature using a 2-fold molar excess of biotin-LC-LC-NHS dissolved in anhydrous DMSO (both ThermoFisher Scientific). Samples were purified on NAP-5 columns, rebuffered in PBS (VWR) and quantified by absorption at 280 nm using a Nanodrop spectrophotometer (ThermoFisher Scientific). Successful biotinylation was validated by flow cytometry using anti-biotin-PE staining.

### Cytotoxicity assays and live-cell imaging

Target cells (2 × 10^4^/well) were co-cultured with CAR^+^ or AdCAR^+^ T cells at indicated effector-to-target (E:T) ratios. For AdCAR assays, biotinylated adapters were added at specified concentrations. Cytotoxicity was monitored as reduction in GFP-positive area using the IncuCyte® S3 Live-Cell Analysis System (Sartorius) and the supplied software (v2019A and v2023A). GFP area was normalized to the first time point. Following co-culture, T cell activation markers were analyzed by flow cytometry and cytokines quantified from supernatants using the MACSPlex™ Cytokine 12 Kit (Miltenyi Biotec). Data were analyzed with MACSPlexInspectoR software (https://www.miltenyibiotec.com/DE-en/resources/tools/macsplex-inspector.html).

### Immunoprecipitation, SDS-PAGE and peptide mass fingerprinting

For each condition, 1 × 10^7^ cells were lysed for 30 min on ice in Ecosurf™ EH-9 lysis buffer (Miltenyi Biotec) containing Halt™ Protease Inhibitor Cocktail (ThermoFisher Scientific). Immunoprecipitation was performed using antibodies with Protein A or biotinylated antibodies with anti-biotin microbeads, followed by μMACS™ Protein A/G separation (all Miltenyi Biotec), according to the manufacturer recommendations. Immunoprecipitates were reduced with 10 mM DTT for 5 min at 95°C and alkylated with 20 mM iodoacetamide (IAA) for 30 min at room temperature. Samples were separated on Novex™ WedgeWell™ 4-20% Tris-Glycine gels using the XCell SureLock™ Mini-Cell system with Tris-Glycine SDS running buffer (all ThermoFisher Scientific). Gels were stained with Bio-Safe™ Coomassie G-250 (Bio-Rad) for 1h and destained overnight under agitation. Bands of interest were excised, destained and digested overnight with trypsin (20 µg/mL) in 50 mM NH4HCO3. Peptides were analyzed by peptide mass fingerprinting (PMF) using an Orbitrap Fusion Lumos Tribrid Mass Spectrometer (ThermoFisher Scientific). Proteins were identified by database searching against the UniProt database.

### Affinity maturation

*In silico* affinity maturation was performed using the Efficient Evolution of Human Antibodies from Protein Language Models package (v0.1)^24^. For yeast surface display (YSD), WT and mutated CLA scFv OligoPools (IDT) were cloned into pYSDM2.0 by yeast recombinational cloning, as previously described^25^. Libraries containing 1-6 random mutations per clone, restricted to the CDR regions, were transformed into electrocompetent *S. cerevisiae* EBY100 as previously described^26^, resulting in libraries in which each yeast clone displays a single scFv variant. Following induction in SGCAA medium^22^, scFvs were labelled via HA-tag APC staining and non-expressing clones depleted using anti-APC microbeads and LD/LS columns (Miltenyi Biotec). Negative selection was performed by incubating yeast with CLA-deficient HL60 FUT7 KO cells at a 1:10 target-to-yeast ratio, followed by sorting using a MACSQuant® Tyto® sorter (Miltenyi Biotec). Positive selection was conducted the following week under similar conditions using WT HL60 cells. To ensure complete CLA depletion in HL60 FUT7 KO cells and robust CLA expression in HL60 WT cells, cells were respectively depleted or enriched using anti-CLA antibodies and LD/LS columns (Miltenyi Biotec) immediately prior to co-culture. After selection, single clones were isolated on agar plates, induced, and screened in 96-well format against WT or FUT7 KO HL60 cells. Clones were analyzed by flow cytometry and ranked according to WT/KO binding ratios and sequenced (Eurofins) to identify mutations.

### Multiplex RT-qPCR

HL60, OCI-AML2 and T cells were cultured at varying densities for 5 days before RNA extraction using the RNeasy® Mini Kit (Qiagen). RNA integrity and concentration were assessed using Qubit™ RNA HS kit (ThermoFisher Scientific) and RNA ScreenTape 4150 TapeStation assay (Agilent Technologies). Multiplex RT-qPCR was performed using the QuantiTect Multiplex RT-PCR NoROX Kit (Qiagen) on a CFX96 instrument (Bio-Rad) with 50 ng RNA per reaction. Expressions of FUT7, B4GALT1, GCNT1, FUT4, ST3GAL4 and B3GNT3 were quantified using IDT primer/probe mixes with HPRT1 as housekeeping gene. Thermal cycling conditions consisted of reverse transcription at 50°C for 20 min, initial activation at 95°C for 15 min, followed by 50 cycles of 94°C for 45s and 60°C for 75s. Relative gene expression was calculated by the ΔΔCt method, where ΔCt = Ct(gene of interest) − Ct(HPRT1) and ΔΔCt = ΔCt(sample) − ΔCt(control). Fold changes in gene expression were calculated as 2^(−ΔΔCt).

### Glycan array screening

A 300-glycan array was performed by CD BioGlyco using HECA-452 and REA1101 CLA antibodies together with generated CLA scFv-Fc binders. CD66c antibody served as background control. Samples were tested at 0.03 mg/mL. Positive control signal intensities were used for data normalization after background subtraction. After normalization and background subtraction performed by CD BioGlyco, log10 fold changes relative to CD66c controls were calculated and z-score normalized. Glycans with z-scores >3 were selected. Raw data and detailed analyses can be downloaded in the supplementary materials.

## 3 Results

### CLA is highly expressed in PDAC but previously reported direct CLA CARs lack functionality

CLA has been described as a glycosylated sialyl-Lewis X epitope mainly carried on CD162/PSGL-1 (figure 1.A). It is highly expressed on multiple PDAC cell lines (BxPC3, AsPC1, Capan2, PanCa0201) and on patient tissues (figures 1.B-C). CLA is also detected on leukemia cell lines (HL60, OCI-AML2), suggesting broader relevance (figure 1.B). On T cells, basal CLA expression increases upon activation (figures 1.B, S2), consistent with prior reports^27^. Healthy tissue array analysis showed no or only minor background CLA expression outside PDAC (figure S3), supporting a favorable safety profile. The only exception was the kidney cortex, where the CLA antibody appeared to bind to an undefined intercellular region. This staining may reflect secreted antigen binding or nonspecific antibody reactivity and warrants further investigation. However, previously reported scFv based anti-CLA CARs were non-functional for unclear reasons^9^. CLA has a prominent expression on activated T cells and previous constructs missed tags for identification of successful surface expression of the CAR, leaving it unclear whether the dysfunction was due to a lack of expression or other reasons. Myc/His tags can help with CAR identification but can interfere with activity and folding. We therefore generated tagged and untagged constructs (figure S4.A), but all failed to induce killing in two PDAC models (figures 1.D, S4.B).

**Figure 1.**
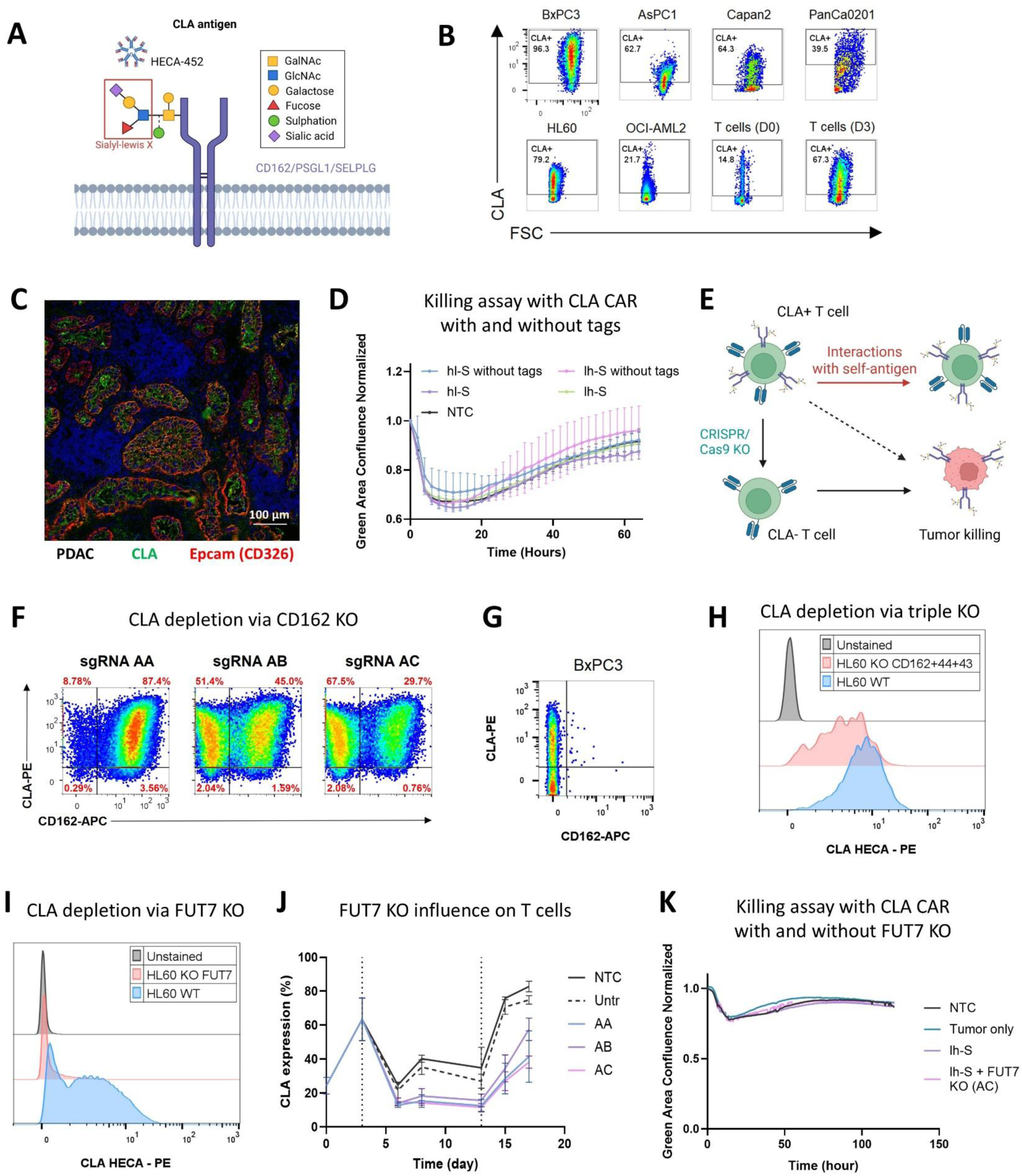
CLA expression and functionality of CLA CAR constructs following different CLA knockout strategies. **A)** Schematic of the CLA antigen, a glycosylated epitope containing sialyl-Lewis X, previously described on CD162/PSGL1/SELPLG and recognized by HECA-452 IgM. Created with BioRender.com. **B)** Flow cytometry analysis of CLA expressions in PDAC cell lines (BxPC3, AsPC1, Capan2, PanCa0201), leukemia cell lines (HL60, OCI-AML2) and T cells at day 0 post-isolation (D0) and day 3 post-activation (D3). Representative of a technical duplicate shown. Gating strategy and controls are provided in figure S1.B. **C)** Cyclic immunofluorescence (IF) of representative human PDAC tissue showing co-expression of CLA and cytokeratin-positive tumor cells. Image is representative of two cyclic IF runs from two different PDAC specimens. **D)** Killing curves showing GFP-based PanCa0201 tumor cell confluence, normalized to the first time point, for hl-S and lh-S CLA CAR constructs with or without Myc and His tags at an E:T ratio of 10:1, compared with NTC. Error bars represent the mean of two independent donors. **E)** Schematic of potential interactions with self-antigen caused by CLA expression on activated T cells and its prevention through CRISPR/Cas9-mediated KO. Created with BioRender.com. **F)** Flow cytometry analysis of CLA depletion in primary activated T cells following CD162 KO using sgRNAs AA, AB and AC. **G)** Flow cytometry analysis of CLA and CD162 expression in BxPC3 cells. Representative of a technical duplicate shown. **H)** Flow cytometry analysis of CLA depletion in HL60 cells following triple CD162/CD43/CD44 KO compared with WT. Gating strategy and controls are provided in figure S5. **I)** Flow cytometry analysis of CLA depletion in HL60 cells following FUT7 KO using sgRNA AC, compared with WT. **J)** CLA expression on peripheral T cells over time. Comparison of FUT7 KO efficiency using sgRNA AA, AB and AC, non-transduced electroporated control (NTC) and untreated control (Untr). Cells were activated with TransAct™ on days 0 and 13. **K)** Killing curves showing GFP-based PanCa0203 tumor cell confluence, normalized to the first time point, for lh-S CLA CAR construct with or without FUT7 KO (sgRNA AC), at an E:T ratio of 2:1, compared with NTC and tumor-only controls, from one donor.

**Figure 2.**
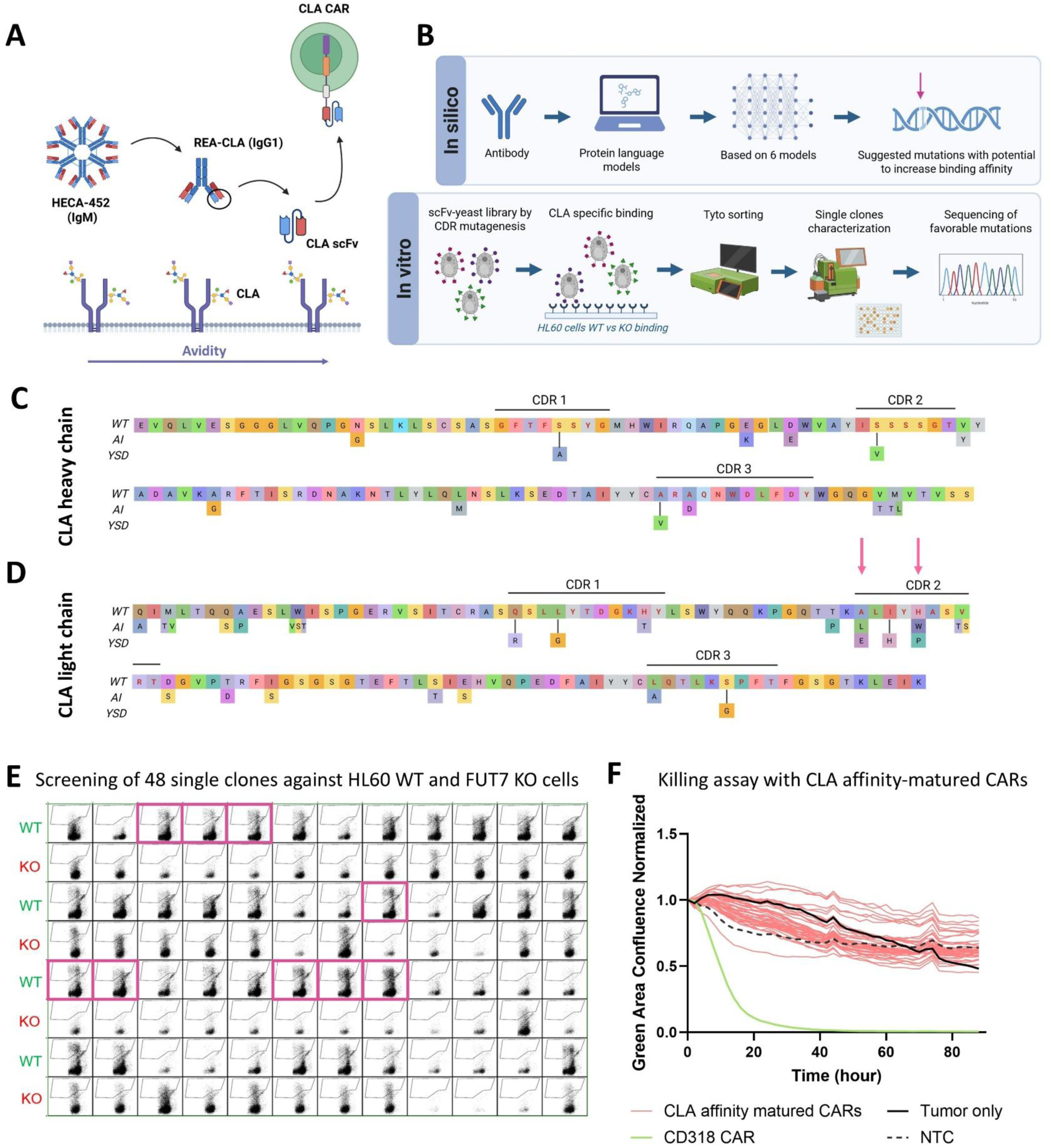
In vitro and in silico CLA affinity maturation to enhance target binding affinity. **A)** Schematic of potential affinity constraints contributing to CAR dysfunction. CLA scFv was derived from the REA IgG1, derived itself from the HECA-452 IgM, reducing binding valency and avidity, potentially impairing affinity. **B)** Affinity maturation workflow combining in silico protein language model prediction and in vitro yeast surface display (YSD). **C-D)** Summary of mutations identified by in silico (AI) and YSD approaches relative to the parental CLA heavy chain **C)** and light chain **D)** sequences. Two mutations in the light-chain CDR2 region (indicated by arrows) were predicted by both methods. The CDRs were defined according to the Chothia numbering scheme. CDR, complementarity-determining region. **A-D)** Created with BioRender.com. **E)** Flow cytometry screening of 48 single clones from a 96-well plate against HL60 WT and FUT7 KO cells. Clones within the pink frame were selected based on the ratio of % binding to WT versus KO cells. **F)** Killing curves showing GFP-based BxPC3 tumor cell confluence, normalized to the first time point, for 30 affinity-matured CLA CAR constructs at an E:T ratio of 2:1, compared with NTC, CD318 CAR and tumor-only controls. Data represents the mean of two independent donors.

### FUT7-mediated CLA depletion does not restore CAR function

Because activated T cells express CLA, we hypothesized that reduced CAR activity was due to cis-or trans-interactions with the self-antigen (figure 1.E)^28^. Since CLA is a glycosylated epitope, direct KO is challenging. We first targeted CD162, the main proposed backbone^11^. Although two sgRNAs (AB, AC) efficiently reduced CD162, CLA levels surprisingly remained unchanged, indicating additional carriers (figure 1.F). Consistently, BxPC3 cells strongly expressed CLA despite lacking CD162 (figure 1.G). As CD44 and CD43 have also been reported as CLA carriers^29^, we generated CD162/CD44/CD43 triple KO cells, but CLA decreased only marginally (10% in HL60 and 4% in activated T cells; figures 1.H, S5). Immunoprecipitation followed by SDS-PAGE and PMF analysis failed to identify alternative backbones, likely due to nonspecific antibody binding (figure S6).

We therefore shifted to FUT7, a key enzyme in CLA biosynthesis. FUT7 KO completely abolished CLA and HECA IgM binding in HL60 cells (figure 1.I). In T cells, TransAct™ stimulation (days 0 and 13) strongly increased CLA, while FUT7 KO reduced this upregulation by ~50% versus controls (figure 1.J). Among the three sgRNAs tested (AA, AB and AC), sgRNA AC showed the highest efficiency and was selected for further experiments (figure S7.A). Interestingly, CLA regulation differed between subsets, with more sustained expression in CD8^+^ cells (figure S7.B-C). In CD4^+^ cells, CLA declined similarly in untreated and KO conditions, whereas CD8^+^ cells showed a slower decrease, indicating more dynamic regulation in CD4^+^ cells. Given that FUT7 may affect broader T cell biology, we further assessed the impact of KO. FUT7 KO did not significantly alter proliferation, activation marker expression (CD69, CD137, CD25, PD1) or differentiation into Tn, Tscm, Tcm, Tem and Teff subsets (figures S7.D-I). Despite efficient CLA depletion, combining FUT7 KO with CLA CARs (e.g. VLVH) did not restore tumor killing, indicating that self-interactions alone do not explain CAR inactivity (figure 1.K).

### Affinity constraints may contribute to dysfunction, yet affinity maturation fails to rescue activity

We next hypothesized insufficient binding affinity. HECA-452 is an IgM antibody, which typically displays low intrinsic affinity but compensates through high avidity due to pentameric structure. CARs use monomeric scFv format, therefore reducing avidity, likely weakening target engagement (figure 2.A). A monomeric recombinant human IgG1 version (REA) confirmed reduced staining compared to IgM (~10-fold lower mean fluorescence intensity (MFI)), supporting reduced effective binding (figures S8.A-B).

To improve binding to CLA, we pursued affinity maturation of the CAR’s scFvs using complementary *in silico* and *in vitro* approaches (figure 2.B). First, we applied a protein language model-based affinity maturation workflow^24^, which predicts evolutionarily plausible antibody mutations without structural or antigen-specific input. Using the anti-CLA scFv sequence, the model identified 30 candidate mutations across framework and CDR regions (figures 2.C-D), ranked by support from six integrated databases (supplementary table 2). In parallel, yeast surface display (YSD) was performed using HL60 cells instead of recombinant antigen (figure S9.A). Negative selection with FUT7 KO cells removed nonspecific binders, followed by positive selection on WT HL60 (figure S9.B). Sorting yielded 1.41% positive cells, which is within the expected range for such selections (figure S9.C). 96 single clones were expanded and re-screened on WT and FUT7 KO cells. Binding specificity was quantified based on the ratio of % binding to WT versus KO cells, enabling selection of target cell specific binders (figure 2.E). Twelve clones were selected showing ≥2.5-fold higher binding signal to WT cells based on frequency. Two clones were confirmed by sequencing, D10 and C2 (with a truncated heavy chain), containing six and three mutations, respectively (figures 2.C-D, S9.D; supplementary table 2). For functional validation, we selected *in silico* mutations supported by ≥4 models, together with YSD-derived mutation combinations (supplementary table 3). However, screening of the 40 selected candidates after affinity-maturation on BxPC3 cells failed to restore cytotoxicity (figure 2.F).

### Functional CLA targeting is achieved via AdCAR using biotinylated scFv-Fc adapters

We next suspected CAR misfolding as a potential cause for CAR dysfunctionality. Indeed, analysis of CAR expression revealed discordant detection of the two encoded tags relative to the LNGFR transduction marker. Myc expression matched LNGFR (~60%), while His-tag was reduced (~24%), suggesting structural masking (figures 3.A-B). To overcome potential folding limitations which might result in affinity reduction, we switched to the AdCAR platform. In this system, biotinylated adapter molecules bind tumor cells and are recognized by anti-biotin AdCAR T cells (figure 3.C)^23^. We generated biotinylated CLA scFv-Fc adapters (VHVL/VLVH). Compared with the original CAR format, scFv-Fc adapters are likely to reduce misfolding risk and present similar avidity through Fc-mediated dimerization, resulting in two binding sites per molecule. Binding analysis on OCI-AML2 and BxPC3 cells showed similar affinities for VHVL and VLVH adapters, comparable to the validated CD66c REA control^30^, and higher binding on BxPC3 cells. Both scFv-Fc adapters exhibited an approximately 100-fold lower EC50 value compared to CLA REA antibody, indicating a significantly higher apparent affinity (figure S10). Both adapters induced dose-dependent cytotoxicity of BxPC3 cells, with near-complete tumor clearance at high VLVH dose (figures 3.D-E). The CLA REA control also exhibited strong, concentration-dependent killing, with the highest activity observed at 10 nM adapter (figure 3.F). The CD66c positive control showed the strongest activity (figure 3.G). Results were reproduced in a second donor, showing similar overall trends despite less distinct dose separation (figure S11).

**Figure 3.**
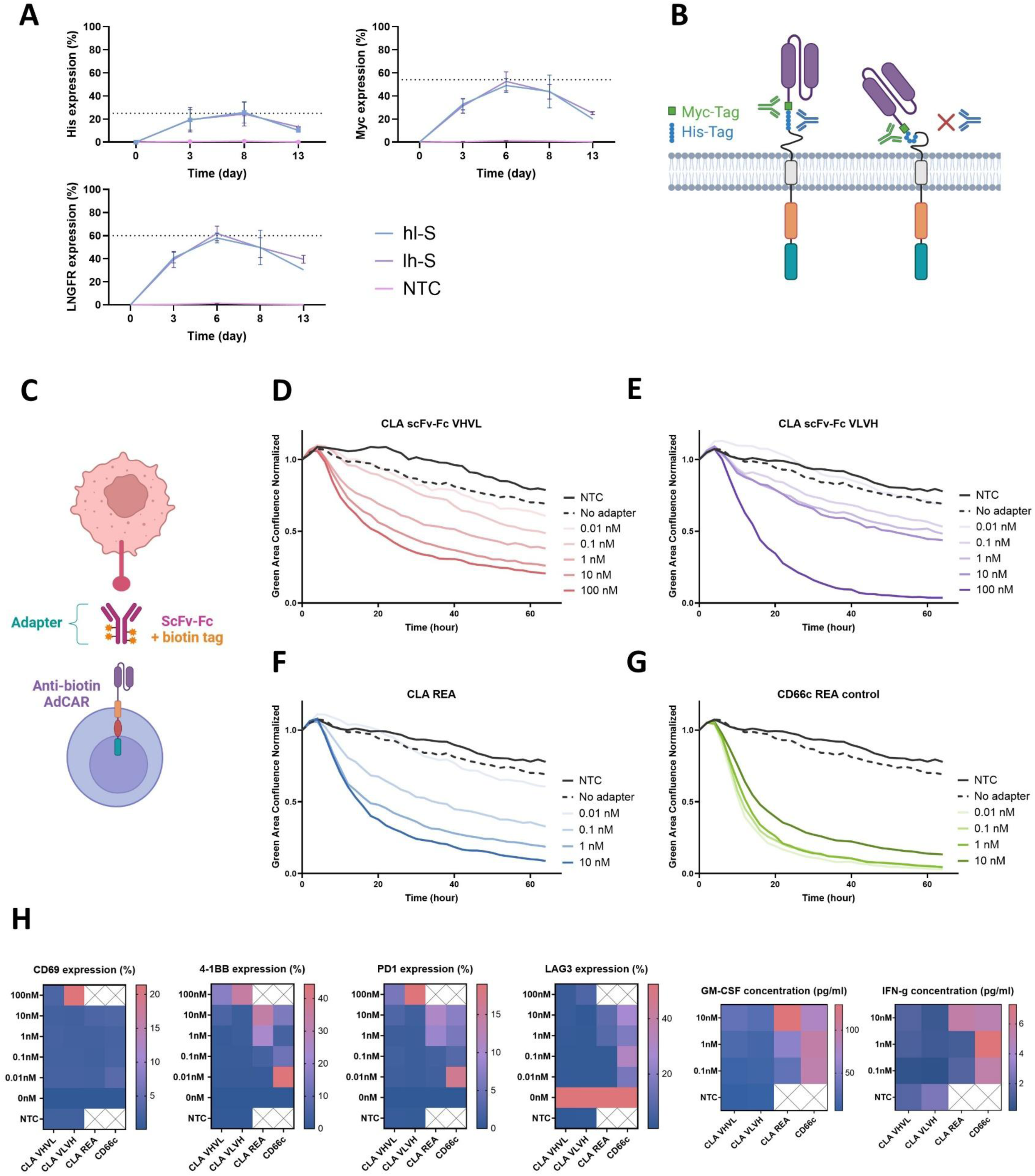
Functional characterization of CLA CARs using biotinylated CLA scFv-Fc adapter molecules in the AdCAR system. **A)** Comparison of CAR expression of hl-S and lh-S CLA CAR T cells over time, detected by His-tag, Myc-tag or LNGFR staining, compared with NTC. Error bars represent the mean of two independent donors. **B)** Schematic of tagged CLA CAR constructs at the cell surface. Left: correctly folded CAR with detectable Myc and His tags. Right: potential CAR misfolding or improper hinge configuration leading to masking of the His tag and loss of anti-His antibody detection. Created with BioRender.com. **C)** Schematic of the AdCAR system, which separates antigen recognition from CAR activation and induces T cell activation only in the presence of specific adapter molecules (e.g., biotinylated scFv-Fc). Created with BioRender.com. **D-G)** Killing curves showing GFP-based BxPC3 tumor cell confluence, normalized to the first time point, for CLA AdCAR constructs at an E:T ratio of 2:1 using a CLA scFv-Fc VHVL adapter **D)**, CLA scFv-Fc VLVH adapter **E)**, CLA REA adapter **F)**, or CD66c REA adapter **G)**, compared with non-transduced T cells (NTC) and AdCAR without adapter (no-adapter) controls, from one donor. Biotinylated adapters were tested at concentrations ranging from 0.01-100 nM for scFv-Fc adapters and from 0.01-10 nM for REA controls. **H)** Heatmaps summarizing CD69, 4-1BB, PD-1 and LAG-3 expression (%) as well as GM-CSF and IFN-γ secretion (pg/mL) across adapter conditions.

Activation correlated with cytotoxicity: CD69, 4-1BB and PD-1 peaked at high VLVH/low CD66c doses, while LAG-3 was highest without adapter as CAR T cells remained inactive (figure 3.H). GM-CSF and IFN-γ production were strongest with CLA REA and CD66c, consistent with their potent killing activity. Interestingly, despite strong cytotoxicity, scFv-Fc adapters induced lower cytokine and activation levels than controls except at highest dose. Overall, CLA mediates effective CAR killing in AdCAR format, with limited fratricide impact.

### CLA variability is density-dependent but remains mechanistically unresolved

We observed pronounced variability in CLA expression across days (figure 4.A). Variation was independent of FBS concentration and did not correlate with tested stress conditions, except for a CLA decrease under constant shaking (figures S12.A-C). Instead, a consistent inverse correlation with cell density was observed across HL60, T cells and OCI-AML2 (figures 4.B, S12.D-E). We next investigated the underlying regulatory factors driving this density-dependent downregulation.

**Figure 4.**
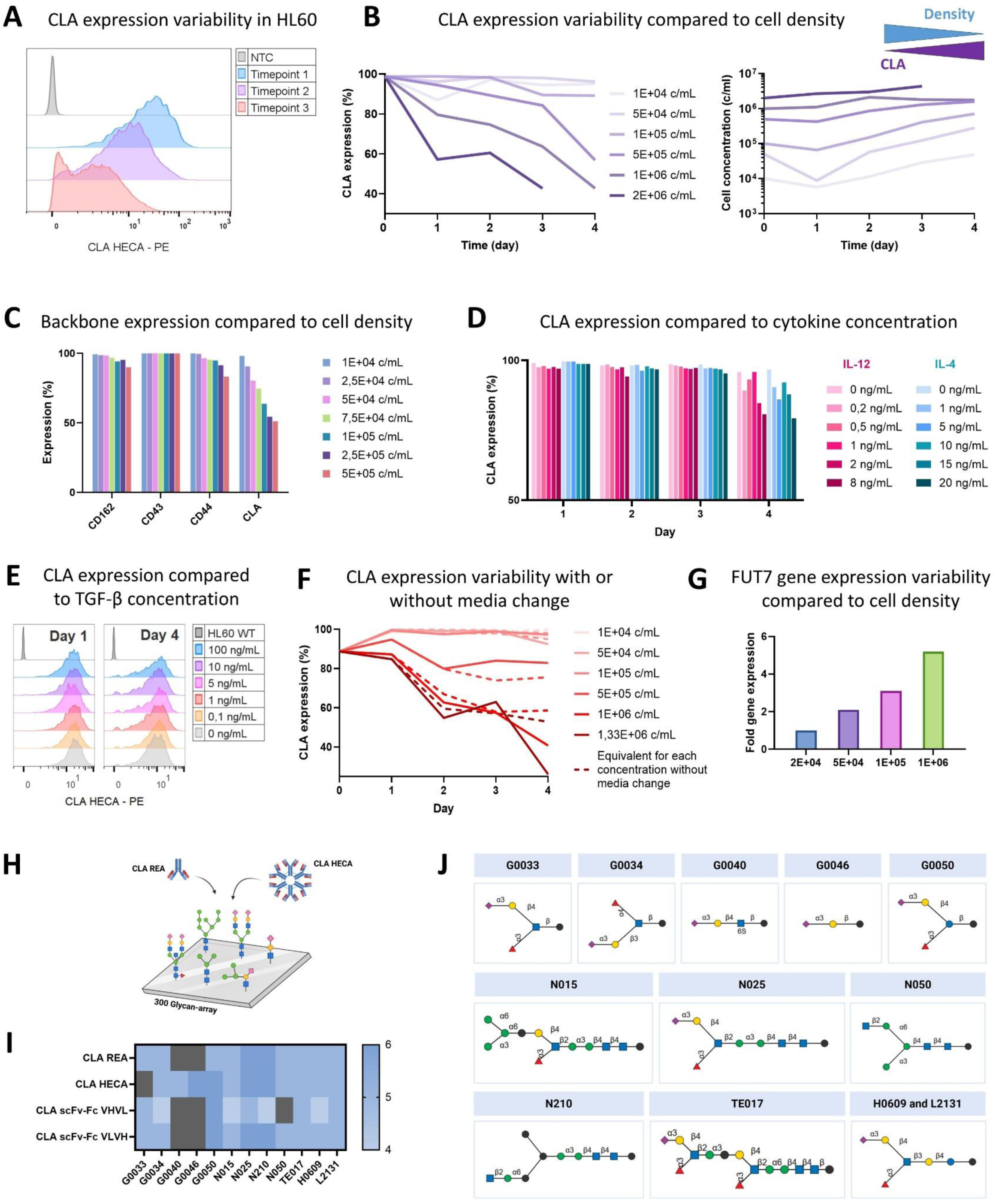
Investigation of density-dependent CLA variability and its underlying regulatory mechanisms. **A)** Flow cytometry analysis of CLA expression variability in HL60 cells at different time points during culture. **B)** Flow cytometry analysis of CLA expression (%) (left) and cell concentration (cells/mL) (right) over time in HL60 cells cultured at starting densities ranging from 1 × 10^4^ to 2 × 10^6^ cells/mL. Representative of a technical duplicate shown. **C)** Flow cytometry analysis of CD162, CD43 and CD44 expression across cell densities (1 × 10^4^ to 5 × 10^5^ cells/mL) compared with CLA expression in HL60. **D)** Flow cytometry analysis of CLA expression over time in HL60 cells cultured with varying IL-12 (0-8 ng/mL) and IL-4 (0-20 ng/mL) concentrations. **E)** Flow cytometry analysis of CLA expression on day 1 and day 4 of HL60 cells cultured with varying TGF-β concentrations (0-100 ng/mL). **F)** Flow cytometry analysis of CLA expression over time in HL60 cells cultured at different cell densities (1 × 10^4^ to 1.33 × 10^6^ cells/mL). From day 1 onward, half of the cultures were maintained unchanged, while the remaining cultures received daily medium replacement (dotted lines) to assess the effects of nutrient depletion and metabolite accumulation. **G)** Bar plot showing fold change in FUT7 expression measured by qRT-PCR in OCI-AML2 cells after 5 days of culture at different cell densities (2 × 10^4^ to 1 × 10^6^ cells/mL). **H)** Schematic of a 300-glycan array using CLA REA and CLA HECA clones to identify glycan-binding specificities and characterize CLA-associated glyco-structures. Created with BioRender.com. **I)** Heatmap of the 12 top glycan candidates showing significantly increased binding (z-score-based) for one or several CLA binders. Each sample was tested in triplicates. **J)** Structures of glycan components corresponding to the 12 top binding candidates. Structures were generated using the GlycoGlyph Toolkit^62^ based on IUPAC-condensed glycan nomenclature. Full glycan names and plate layout are provided in the supplementary materials. Created with BioRender.com.

We first assessed whether changes in known scaffold or carrier proteins could explain reduced CLA expression. Minor decreases in CD162 and CD44 were observed, but these were insufficient to account for the ~50% reduction in CLA at high density (figure 4.C), consistent with previous evidence that CLA is not restricted to these backbones. We next evaluated cytokine-mediated regulation, as increasing cell density alters the cytokine milieu. IL-12 has been reported to promote CLA via skin-homing programs^31^; IL-4 has been reported to suppress enzymes required for CLA synthesis, especially FUT7^32,33^. Finally TGF-β would enhance FUT7 expression in T cells^34^. However, cytokine stimulation induced only a mild, transient CLA decrease on day 4 under high-dose IL-12 and IL-4, with no effect of TGF-β, partially contradicting prior reports (figures 4.D-E). We also tested whether nutrient depletion or metabolite accumulation at high density contributed to reduced CLA. Daily media exchange experiments showed only minimal effects, with no rescue of CLA levels (figure 4.F). Expression analysis of key enzymes involved in CLA biosynthesis (FUT7, B4GALT1, GCNT1, FUT4, ST3GAL4 and B3GNT3^15,35^) showed no consistent downregulation accompanying CLA loss (figure S13). In some cases (e.g., FUT7 in OCI-AML2), expression even slightly increased at higher density (figure 4.G). GCNT1 and B3GNT3 were not reliably detected. Finally, given that CLA is an adhesion-related glyco-epitope and adhesion molecules are often density-regulated, we assessed multiple adhesion markers but observed no correlation with CLA reduction (figure S14). In summary, CLA is consistently downregulated at high cell density across cell types, but the underlying mechanism remains unresolved.

### Broad and heterogeneous glycan recognition by CLA binders challenge the current definition of CLA

As previously noted, contradictory findings in the literature and our study suggest that CLA may be more complex than originally described, and that antibody specificity may be limited. We therefore profiled glycan binding using a 300-glycan array with REA, HECA, and scFv-Fc binders to define CLA-recognized structures (figure 4.H). The canonical CLA structure corresponds to glycan G0033 in this assay. Unexpectedly, all four binders interacted significantly with multiple glycans beyond G0033, and HECA did not significantly bind G0033 (Sialyl Lewis X). In total, 12 glycans were recurrently recognized, although binding profiles varied between binders (figures 4.I-J and S15). Among these, several matched Sialyl Lewis X-related motifs with extended structures (N025, TE017, H0609, L2131) or Sialyl Lewis A-like features (G0034). Others shared partial or truncated motifs (G0050, G0040, G0046), while some (N015, N210) displayed more divergent glycan architectures. Overall, these data suggest either substantial antibody cross-reactivity or that CLA should be defined as a broader glycan family rather than a single epitope.

## 4 Discussion

PDAC remains a devastating disease with limited therapeutic options. However, immunotherapy is opening new avenues, with several promising targets emerging, including CD318, already advancing toward clinical evaluation^36^. Beyond CD318, additional targets such as mesothelin^37^, CEA^38^, claudin 18.2^39^ and HER2^40^ have been explored, but their expression in normal tissues has raised safety concerns and, in some cases, caused severe adverse events, including fatal systemic toxicity in HER2 CAR T cells due to low-level pulmonary expression^41^ and transient colitis in CEA-directed CAR T cells^42^. Within this landscape, we propose CLA as an attractive PDAC target due to its restricted expression in tumor tissue and selected leukocyte subsets^9^. Importantly, we confirmed the absence of detectable expression in healthy tissues, reinforcing its therapeutic interest. The only exception was the kidney cortex, where the observed staining remains unexplained and may represent either secreted antigen binding or nonspecific antibody reactivity, warranting further investigation. Beyond PDAC, CLA has been implicated in lymphoma^43^ and skin inflammatory diseases^44^, highlighting broader biological relevance.

Engineering a functional CLA CAR proved challenging due to the non-standard nature of the antigen. Although uncommon, glycan-targeting CARs have been described, as aberrant glycosylation is a hallmark of cancer^45^. CAR T cells targeting Tn-MUC1 have demonstrated efficacy in leukemia, pancreatic and breast cancer models^46^ and are in phase I evaluation^47^. GD2-directed CARs have shown strong clinical potential across neuroblastoma^48^, glioblastoma^49^, sarcoma and melanoma^50^. Finally, preclinical studies have shown Lewis Y CARs efficacy in prostate cancer models^51^. In PDAC, altered glycosylation, including high-mannose glycans, is prominent, and lectin-based H84T BanLec CAR T cells have shown potent preclinical activity^52,53^. These studies collectively support glyco-epitopes as viable CAR targets, conceptually aligned with CLA.

However, engineering glycan-targeting CARs remains challenging due to difficulties in generating high-affinity and specific binders. Antibody development against glycans is often limited by weak immunogenicity and tolerance to human glyco-structures in murine systems^45^. For CLA, we relied on its natural HECA-452 IgM binder and derived IgG and scFv formats. To improve CAR performance, we tested common failure modes, including hinge optimization or tag removal, but this was insufficient^9^. We therefore investigated self-binding, misfolding and affinity limitations. Self-binding was prevented by FUT7 KO, resulting in CLA depletion on T cells; however, this did not restore the cytotoxic activity of the CLA CAR. Potential misfolding caused by the Myc and His tags was investigated, but tag removal likewise failed to improve killing. To assess the functionality of the scFv itself, scFv-Fc fusion proteins were generated and used in the AdCAR system. Their compatibility with the AdCAR platform confirmed that the scFv remained functional. Although a global folding issue of the complete CAR construct cannot be excluded, the data increasingly pointed toward an affinity-related limitation.

Affinity maturation using *in silico* and YSD approaches identified multiple candidates but failed to generate functional CARs. Nevertheless, we successfully adapted a glycan-compatible YSD workflow using HL60 cells with WT/KO dual selection, broadening the applicability of the method. Alternative approaches such as phage display could offer broader sequence exploration, while yeast display remains superior for fine affinity tuning. However, neither strategy addresses higher-order CAR constraints such as folding, orientation or dimerization. CAR pooling or functional screening approaches (based on CAR activity rather than binder affinity) may better capture these effects in future studies^54^. *In silico* affinity maturation generated multiple mutations but no functional CARs, likely due to model limitations trained on full antibodies rather than scFvs and lack of structural constraints^24^. Many candidates were germline-like substitutions with uncertain benefit. Emerging models such as BoltzGen^55^, OrthoRep^56^ or RFdiffusion^57^ may overcome these limitations.

Despite excluding individual failure modes, no functional CAR initially emerged, suggesting combined interdependent constraints rather than a single dominant cause. We therefore transitioned to an AdCAR strategy, which overcomes several limitations of direct CARs such as dependence on antigen density and epitope geometry. AdCARs exploit multivalent adapter-mediated interactions, enhancing avidity and flexibility, while scFv-Fc adapters improve valency and stability, potentially reducing misfolding. Using biotinylated CLA scFv-Fc adapters, we achieved robust dose-dependent cytotoxicity, validating CLA as a functional PDAC target for the first time. Importantly, efficient killing occurred without prior FUT7 KO, suggesting limited self-antigen impact. Another noteworthy observation was that the scFv-Fc format required approximately 10-fold higher adapter concentrations than the REA format to achieve comparable cytotoxicity, despite that the REA had lower titrated binding performance. These stochiometric differences could be explainable by the multitude of parameters playing into the functionality of the AdCAR system. While affinities may be different between the different antibody variants, the degree of biotinylation might as well. As we did not investigate the influence of the degree of biotinylation, it is hard to pin down the exact reason for discrepancies in the binding affinity and killing. While this finding warrants further characterization in the context of preclinical development, it did not impair CAR function here.

Beyond engineering aspects, this work also provides new insights into CLA biology. We confirm FUT7 as essential for CLA expression and its knockout does not impair T cell function. In contrast, CD162, CD44 or CD43 KO did not abolish CLA expression, indicating that additional, as yet unidentified, CLA carriers exist. These carriers may include other glycoproteins or, potentially, non-protein molecules such as glycosphingolipids, which have been reported to carry carbohydrate structures related to CLA, including the sialyl Lewis X motif^58^. Proteomic identification was inconclusive, likely due to nonspecific binding or potentially dynamic redistribution of CLA across multiple scaffolds. Glycan arrays revealed large and heterogeneous binding of CLA binders, supporting either differential recognition or a broader CLA glyco-epitope family. The implication of this finding is that our previous analysis of the HECA clones may not be entirely accurate, as we show that this binding structure exhibits promiscuous binding properties.

We also identified strong cell density-dependent regulation of CLA across multiple cell types. We performed extensive investigation, including cytokine modulation (IL-12, IL-4, TGF-β), protein expression (backbone and adhesion molecule regulation) or metabolic modulation (nutrient depletion, metabolic product accumulation and enzymes regulation). However, we could not identify a single responsible mechanism. This could suggest a multifactorial or context-dependent regulation that would need further investigation. Future studies could also be extended by performing cell cycle analyses or by examining the expression and activity of enzymes involved in CLA degradation or, more broadly, glycosphingolipid metabolism, as the expression of these enzymes is often influenced by cell density^59–61^.

In conclusion, this study demonstrates that CLA is a viable PDAC CAR target when implemented via AdCAR, overcoming intrinsic limitations of direct CAR design. We further uncover density-dependent regulation and incomplete structural definition of CLA. While further work is needed to fully resolve its molecular identity and regulatory network, CLA represents a promising target not only for cancer immunotherapy but potentially also for inflammatory and dermatological diseases.

## 5 Data Availability Statement

The datasets used and/or analyzed in this paper are available from the corresponding authors on reasonable request. The glycol-array raw data and analyzed results are available in supplementary files.

## 6 Ethics statement

PBMCs were isolated from the whole blood of healthy anonymous volunteers at Miltenyi Biotec, with written informed consent as approved by the local ethics committee of Ärztekammer Nordrhein (2020272). All blood samples were handled following the required ethical and safety procedures.

## 7 Author Contributions

C.D and D.S conceived the study and developed the experimental strategies. C.D performed the staining, killing assays, the triple CD43/CD44/CD162 KO, the CLA affinity maturation, the AdCAR assay, the CLA regulation investigations and the CLA glycan analysis. C.D further performed the data analysis and the manuscript preparation. L.G and D.S performed the CLA direct-CAR designs, the FUT7 and CD162 KO investigations and analysis. D.S performed the cyclic immunofluorescence staining and analysis. K.V supported C.D for the cytotoxicity assay handling and manuscript revision. N.B, M.D and A.F supported C.D for the affinity maturation by yeast display. S.T supported the glyco-array analysis and investigated CLA backbones and glycosylation enzymes. H.H and J.W produced the anti-CLA scFv-Fc molecules. D.S and O.H contributed to supervision. All authors contributed to manuscript revision.

## 8 Funding

This work was supported and funded by Miltenyi Biotec B.V. & Co. KG. The authors gratefully acknowledge the company’s financial and technical support, which enabled the execution of experiments described herein.

## 9 Acknowledgments

The authors thank the other members of Binder Generation team at Miltenyi for their support for the affinity maturation. We further thank Felix Heider, Sarah Kribben and Anna-Lena Zinn for KO support; Svenja Oberbörsch for assistance with SDS-PAGE analysis; Ramona Braun and Jens Hellmer for PMF mass spectrometry identification; Lukas Kiefer and Michael Wagner for advice on data analysis; Martina Hilger for CLA REA and HECA antibody titration; Martin Helm for support with *in silico* affinity maturation; Vera Dittmer for the MACSima run with healthy tissues; Julia Schnell and Matthias Bernhard Wahl for assistance with RT-qPCR assays; Lena Willnow for support with AdCAR biotinylation; Nele Knelangen and Ulrika Bader for providing the AdCAR lentivectors; Aleksej Frolov and Camille Métais for glycan array analysis; Anniek Timmermans and Dave Huang for their contributions to the project as master’s students.

The authors would like to acknowledge the use of BioRender.com with a valid license in the creation of figure 1.A (https://BioRender.com/eo6xy0w), figure 1.E (https://BioRender.com/6kqreqj), figure 2.A (https://BioRender.com/h4t5yje), figure 2.B (https://BioRender.com/5o990sg), figures 2.C-D (https://BioRender.com/n2exp97), figure 3.B (https://BioRender.com/3t91cqz), figure 3.C (https://BioRender.com/dhjbnht), figure 4.H (https://BioRender.com/6o4ushi), figure 4.J (https://BioRender.com/viv0xt3), figure S1 (https://BioRender.com/q6dp5qj), figure S4 (https://BioRender.com/e0339e1), figure S5 (https://BioRender.com/zxq7s89), figure S9.A (https://BioRender.com/moo4sq3), figure S9.B (https://BioRender.com/nwlei0b) and figure S12 (https://BioRender.com/bdws7bc), all licensed under CC BY 4.0.

## 10 Conflict of Interest

C.D, K.V, N.B, M.D, A.F, J.W, S.T, H.H, O.H and D.S are employees of Miltenyi Biotec B.V. & Co. KG. L.G was an employee of Miltenyi Biotec B.V. & Co. KG at the time the experiments were performed. All other authors declare no competing interests.

## 11 Generative AI statement

The author(s) declare that ChatGPT (GPT-5.3, OpenAI) was used in the revision of this manuscript, to improve the readability and language. After using this tool, the authors reviewed the content critically and thoroughly, edited it wherever needed, and take full responsibility for the content of the publication. All scientific content, interpretations and conclusions were generated and verified by the authors.

## Supplementary tables

**Supplementary table 1.** List of Miltenyi Biotec antibodies used for flow cytometric analysis, cyclic immunofluorescence and CLA carrier protein identification.

| Target | Clone | Conjugate |
| --- | --- | --- |
| AdCAR detection reagent | na | PE |
| Anti-biotin | REA746, human | PE |
| Anti-HA | REA1288, human | APC |
| Anti-His | GG11-8F3.5.1, mouse | FITC and PE |
| Anti-Myc | SH1-26E7.1.3, mouse | FITC |
| CLA HECA-452 | HECA-452, rat IgM | PE and unconjugated and biotinylated |
| CLA REA | REA1101, human IgG1 | PE and unconjugated |
| CD2 | REA972 | PE-Vio770 |
| CD4 | REA623, human | VioGreen |
| CD8 | REA734, human | VioBlue |
| CD11a | REA378, human | PE |
| CD25 | REA570, human | PE-Vio770 |
| CD43 | REA833, human | VioGreen |
| CD44 | REA690, human | VioBlue |
| CD45 | REA747, human | Unconjugated and biotinylated |
| CD45RA | REA562, human | VioBlue |
| CD45RO | REA611, human | APC |
| CD49d | REA545, human | PE |
| CD54 (ICAM-1) | REA266, human | APC |
| CD58 (LFA-3) | REA1098, human | PE-Vio770 |
| CD62L | REA828, human | PE-Vio770 |
| CD66c | REA414, human | Unconjugated and biotinylated |
| CD69 | REA824, human | APC-Vio770 |
| CD95 | REA738, human | PE |
| CD106 (VCAM-1) | REA269, human | PE-Vio770 |
| CD137 (4-1BB) | REA765, human | APC |
| CD162 (PSGL1) | REA319, human | APC, unconjugated and biotinylated |
| CD183 (CXCR3) | REA232, human | PE |
| CD184 (CXCR4) | REA649, human | APC-Vio770 |
| CD197 (CCR7) | REA108, human | APC |
| CD223 (LAG3) | REA351, human | VioBlue |
| CD271 (LNGFR) | REA844, human | APC |
| CD279 (PD1) | REA1165, human | PE-Vio770 |
| CD326 (Epcam) | REA764, human | FITC |

**Supplementary table 2.**
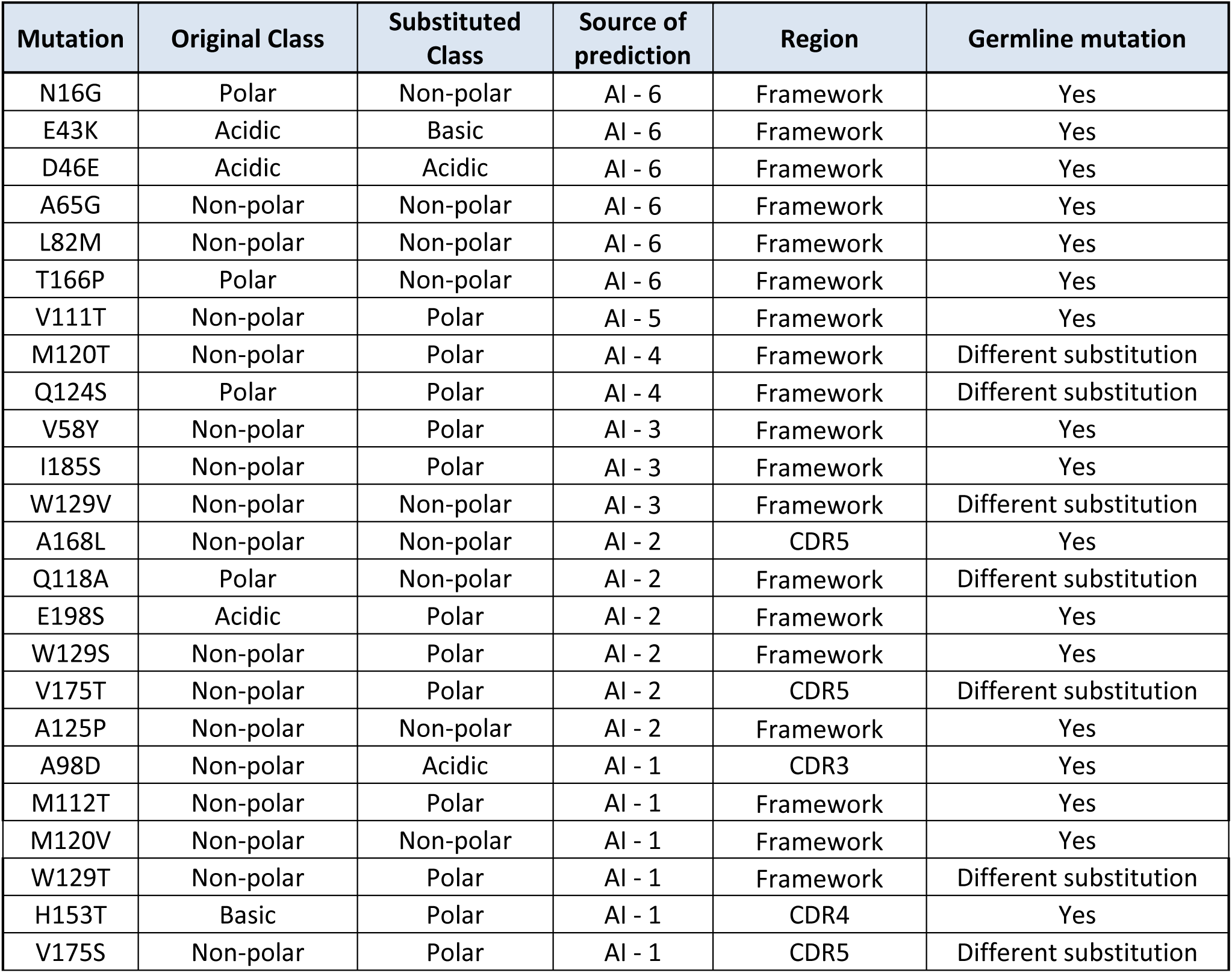
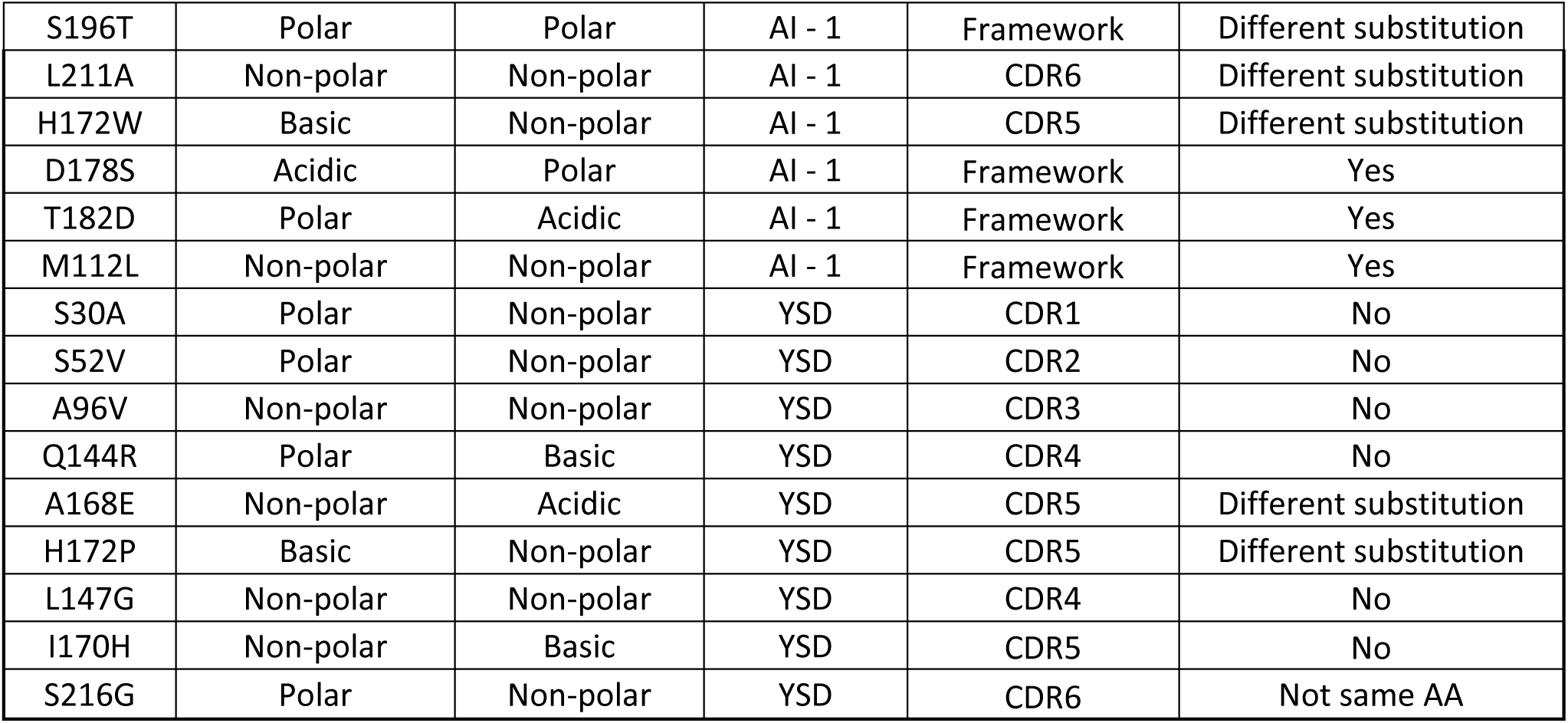
List of mutations identified through in silico and in vitro affinity maturation. For each mutation, the original amino acid class and the substituted amino acid class are indicated. The source of the prediction is specified as either in silico (AI), including the number of models supporting the mutation (1 to 6), or in vitro using yeast surface display (YSD). The location of each mutation is annotated as belonging to either a framework region or a complementarity-determining region (CDR). Information on whether each mutation corresponds to a germline residue (yes/no) is provided, according to IMGT/V-QUEST. Mutations at germline positions but involving a different amino acid labelled as “different substitution”.

**Supplementary table 3.** List of affinity-matured CLA CAR variants containing mutations identified through in silico and in vitro affinity maturation. Due to the large number of predicted mutations and the limited throughput of CAR cloning and functional assays, 40 candidate combinations were selected for testing. All in silico mutations supported by at least four prediction models were included. In addition, selected mutations supported by three, two or one model(s) were incorporated, preferentially within CDR regions, to ensure representation across confidence categories. A CAR containing all the 12 mutations supported by at least three models was also generated. All mutations identified by YSD were tested individually, in combination and in “minus-one” combinations to identify mutations critical for affinity improvement. Because clone C2 contained a truncated heavy chain, all corresponding C2-derived combinations were additionally evaluated in a truncated heavy-chain CLA CAR format.

| Affinity matured CAR | Source of prediction | Mutation(s) | Description |
| --- | --- | --- | --- |
| 1 | AI (*in silico*) | N16G | Supported by 6 models |
| 2 | AI (*in silico*) | E43K | Supported by 6 models |
| 3 | AI (*in silico*) | D46E | Supported by 6 models |
| 4 | AI (*in silico*) | A65G | Supported by 6 models |
| 5 | AI (*in silico*) | L82M | Supported by 6 models |
| 6 | AI (*in silico*) | T166P | Supported by 6 models |
| 7 | AI (*in silico*) | V111T | Supported by 5 models |
| 8 | AI (*in silico*) | M120T | Supported by 4 models |
| 9 | AI (*in silico*) | Q124S | Supported by 4 models |
| 10 | AI (*in silico*) | V58Y | Supported by 3 models |
| 11 | AI (*in silico*) | W129V | Supported by 3 models |
| 12 | AI (*in silico*) | A168L | Supported by 2 models |
| 13 | AI (*in silico*) | V175T | Supported by 2 models |
| 14 | AI (*in silico*) | A98D | Supported by 1 model |
| 15 | AI (*in silico*) | V175S | Supported by 1 model |
| 16 | AI (*in silico*) | L211A | Supported by 1 model |
| 17 | AI (*in silico*) | H172W | Supported by 1 model |
| 18 | AI (*in silico*) | N16G, E43K, D46E, A65G, L82M, T166P, V111T, M120T, Q124S, V58Y, I185S, W129V | All 12 mutations supported by at least 3 models combined |
| 19 | YSD (*in vitro*) | S30A, S52V, A96V, Q144R, A168E, H172P | All 6 mutations supported by clone D10 |
| 20 | YSD (*in vitro*) | S52V, A96V, Q144R, A168E, H172P | 5 mutations supported by clone D10 without S30A |
| 21 | YSD (*in vitro*) | S30A, A96V, Q144R, A168E, H172P | 5 mutations supported by clone D10 without S52V |
| 22 | YSD (*in vitro*) | S30A, S52V, Q144R, A168E, H172P | 5 mutations supported by clone D10 without A96V |
| 23 | YSD (*in vitro*) | S30A, S52V, A96V, A168E, H172P | 5 mutations supported by clone D10 without Q144R |
| 24 | YSD (*in vitro*) | S30A, S52V, A96V, Q144R, H172P | 5 mutations supported by clone D10 without A168E |
| 25 | YSD (*in vitro*) | S30A, S52V, A96V, Q144R, A168E | 5 mutations supported by clone D10 without H172P |
| 26 | YSD (*in vitro*) | L147G, I170H, S216G | All 3 mutations supported by clone C02 |
| 27 | YSD (*in vitro*) | I170H, S216G | 2 mutations supported by clone C02 except L147G |
| 28 | YSD (*in vitro*) | L147G, S216G | 3 mutations supported by clone C02 except I170H |
| 29 | YSD (*in vitro*) | L147G, I170H | 4 mutations supported by clone C02 except S216G |
| 30 | YSD (*in vitro*) | L147G, I170H, S216G | All 3 mutations supported by clone C02 with the light chain only |
| 31 | YSD (*in vitro*) | S30A | Single mutation supported by clone D10 |
| 32 | YSD (*in vitro*) | S52V | Single mutation supported by clone D10 |
| 33 | YSD (*in vitro*) | A96V | Single mutation supported by clone D10 |
| 34 | YSD (*in vitro*) | Q144R | Single mutation supported by clone D10 |
| 35 | YSD (*in vitro*) | A168E | Single mutation supported by clone D10 |
| 36 | YSD (*in vitro*) | H172P | Single mutation supported by clone D10 |
| 37 | YSD (*in vitro*) | L147G | Single mutation supported by clone C2 |
| 38 | YSD (*in vitro*) | I170H | Single mutation supported by clone C2 |
| 39 | YSD (*in vitro*) | S216G | Single mutation supported by clone C2 |
| 40 | na | na | CLA parental clone (WT) |

## Supplementary figures

**Supplementary figure S1.**
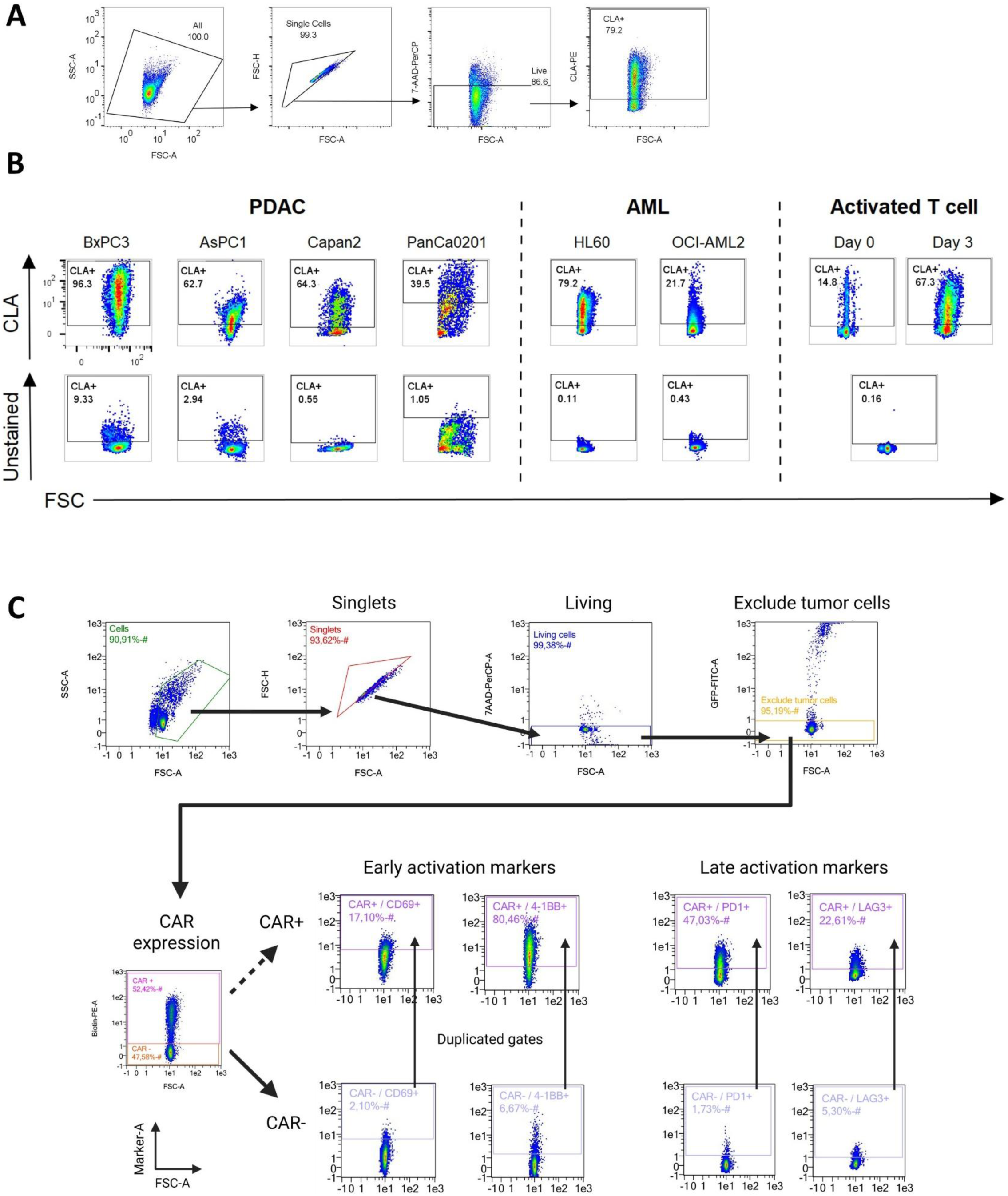
Flow cytometry gating strategies. **A)** Gating strategy for single-marker analysis (e.g., CLA). Initial gating steps excluded debris (FSC-A vs SSC-A), doublets (FSC-A vs FSC-H) and dead cells (FSC-A vs 7AAD/PerCP-A). The marker of interest was gated relative to unstained control (NTC). **B)** Gating strategy for CLA analysis in PDAC and leukemia cell lines, as well as T cells, compared with their respective unstained controls (NTC). **C)** Gating strategy for activation marker analysis following CLA AdCAR killing assays. Initial gating steps excluded debris (FSC-A vs SSC-A), doublets (FSC-A vs FSC-H) and dead cells (FSC-A vs 7AAD/PerCP-A). CAR^+^ T cells were identified as biotin-PE^+^. Early activation was assessed by CD69 and 4-1BB expression, and late activation by PD-1 and LAG-3 expression. Gates were defined using CAR^−^ cells and the non-transduced control (NTC), then applied to CAR^+^ cells to ensure unbiased thresholding across T-cell populations. Created with BioRender.com.

**Supplementary figure S2.**
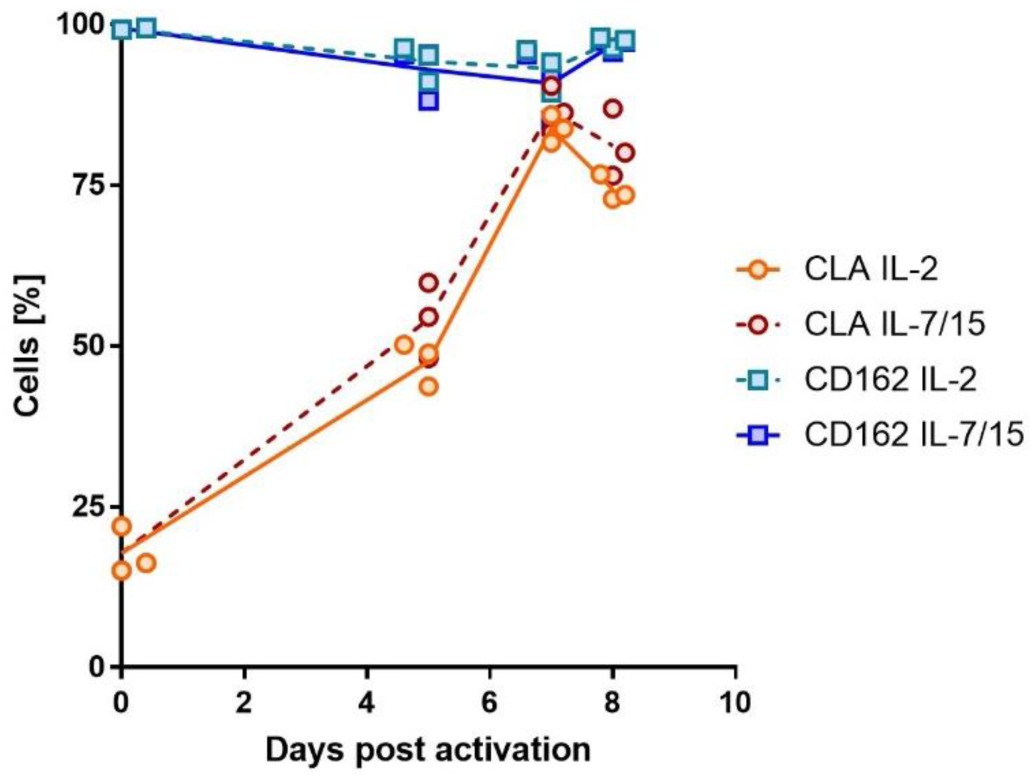
Flow cytometry analysis of CLA and CD162 expression on T cells following activation with TransAct™ over time. Data represents the mean of T cells from three independent donors cultured in medium supplemented with either IL-2 (12.5 ng/mL) or IL-7 and IL-15 (12.5 ng/mL each).

**Supplementary figure S3.**
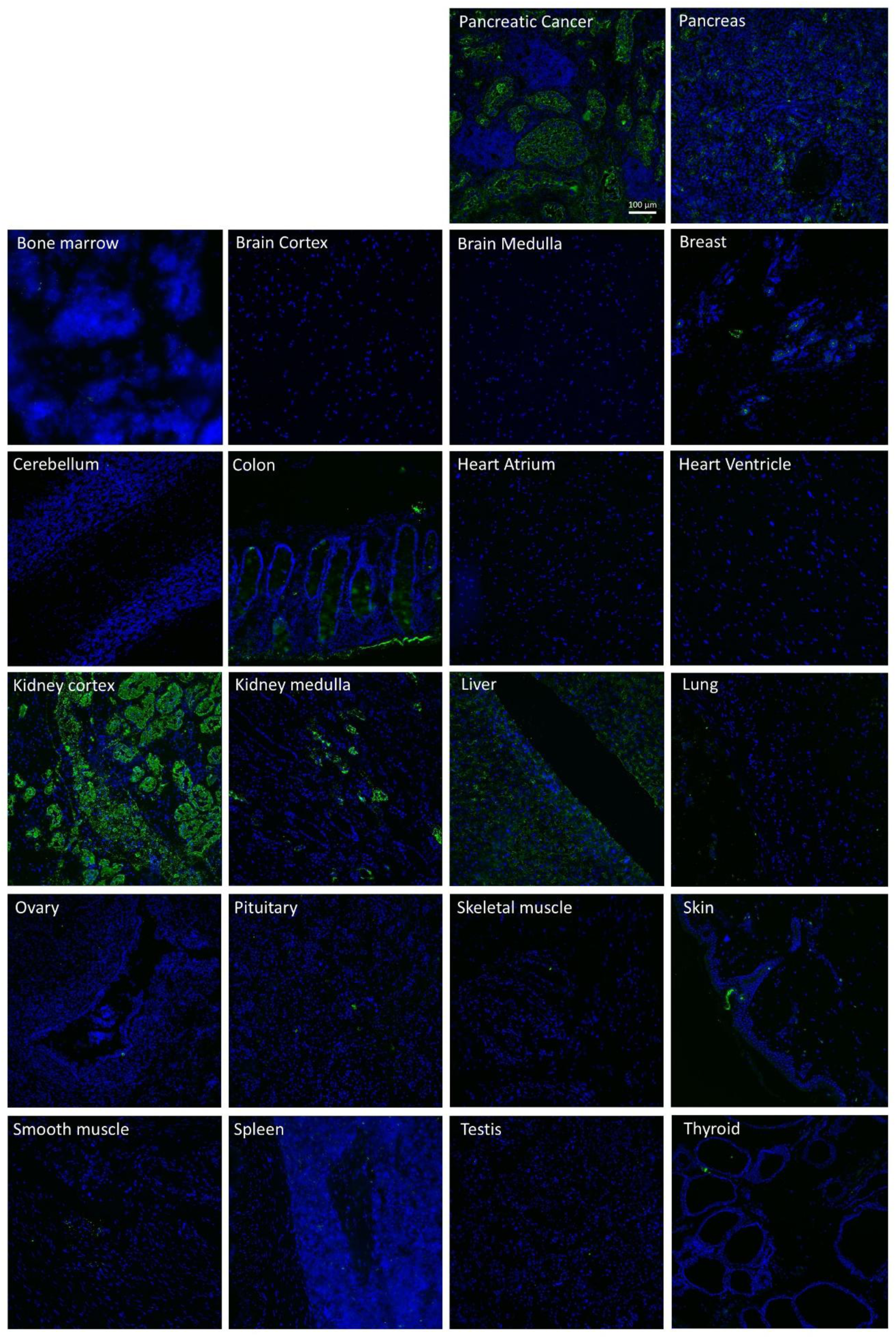
CLA expression on healthy tissues assessed by cyclic immunofluorescence imaging. Representative cyclic immune fluorescence images of several healthy tissues stained with an anti-CLA-PE conjugate (HECA-452). Scale bar = 100 µm. Images are representative of at least two regions of interest from one tissue. Regions of interest were chosen based on manual DAPI and Cytokeratin prestaining and dependency on the respective tissue size.

**Supplementary figure S4.**
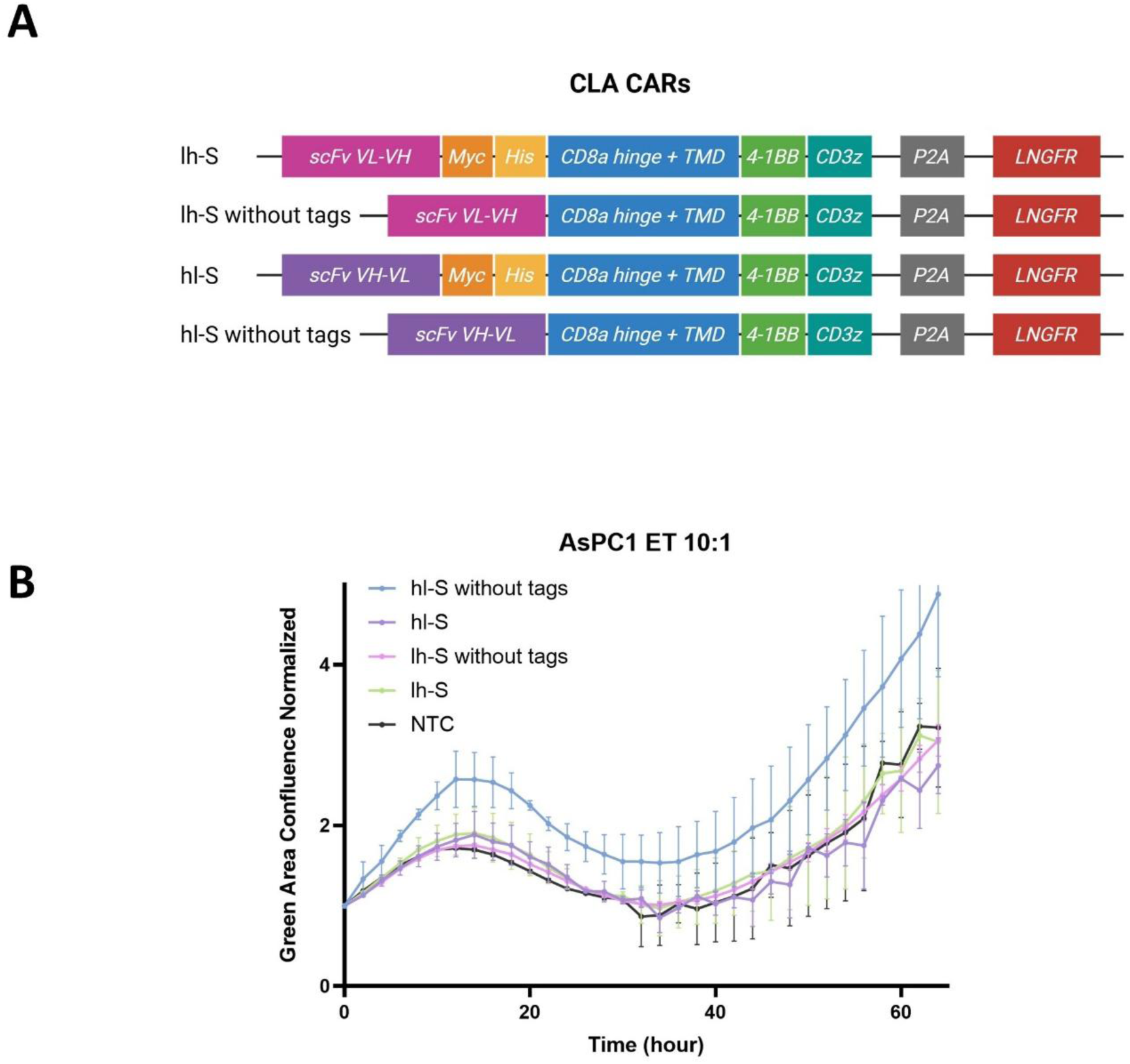
Test of the functionality of the initial CLA CAR designs. **A)** Schematic of the second-generation CLA CAR cassette containing two scFv orientations (VLVH or VHVL, referred to as lh and hl respectively), a short CD8α hinge (S), a transmembrane domain (TMD), a 4-1BB co-stimulatory domain, a CD3ζ signaling domain, and optional Myc and His tags. The cassettes are followed by a Porcine teschovirus-1 2A self-cleaving peptide (P2A) and a low-affinity nerve growth factor receptor (LNGFR) as transduction marker. Created with BioRender.com. **B)** Killing curves showing GFP-based AsPC1 tumor cell confluence, normalized to the first time point, for hl-S and lh-S CLA CAR constructs with or without Myc and His tags at an E:T ratio of 10:1, compared with NTC. Error bars represent the mean of two independent donors.

**Supplementary figure S5.**
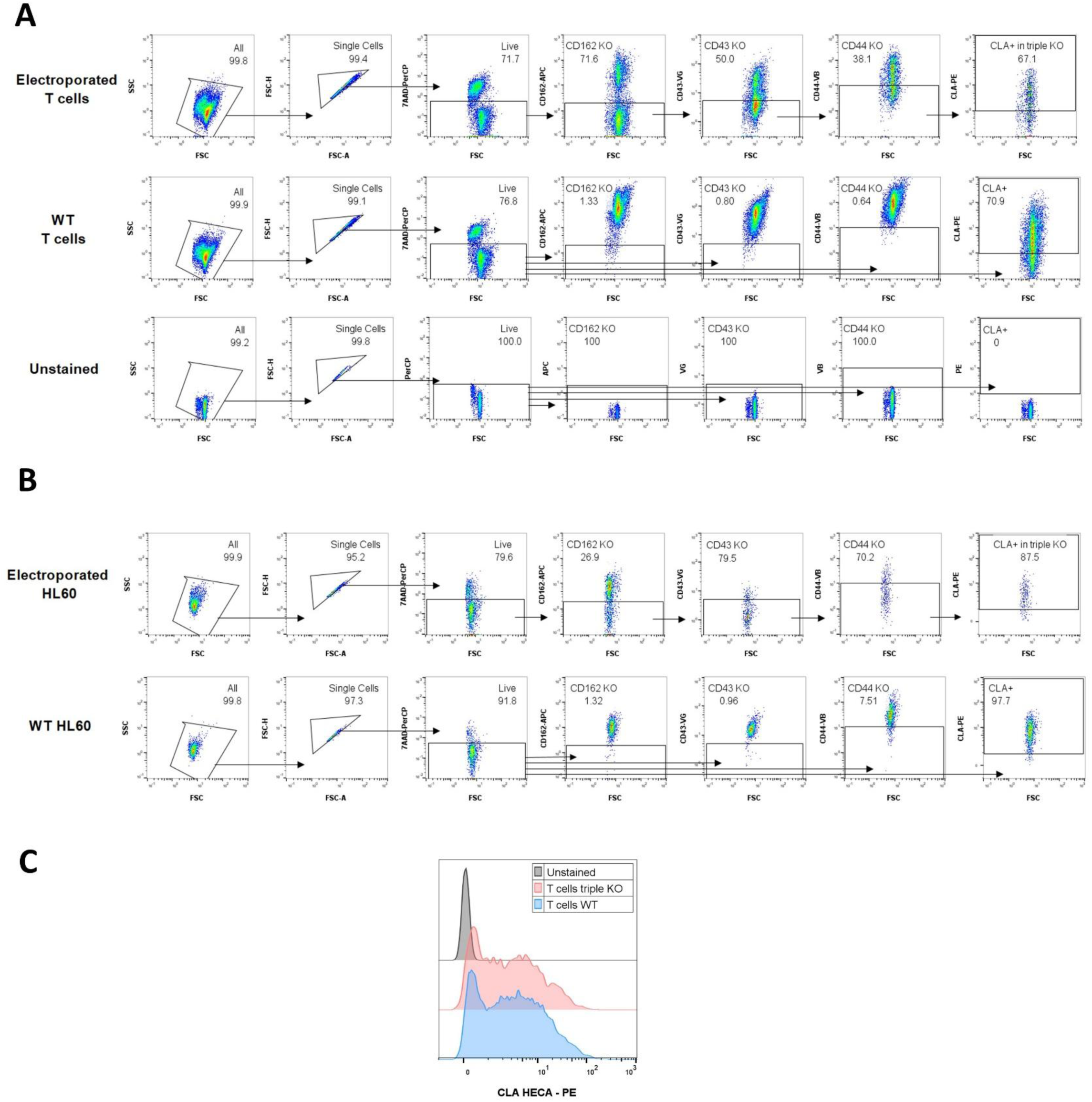
CLA depletion through CRISPR/Cas9-mediated CD44, CD43 and CD162 triple KO in multiple cell lines. **A-B)** Gating strategy for triple KO analysis in T cells **A**) and HL60 cells **B**). Initial gating excluded debris (FSC-A vs SSC-A), doublets (FSC-A vs FSC-H) and dead cells (FSC-A vs 7AAD/PerCP-A). CD162, CD43 and CD44 expression were gated relative to unstained (NTC) and the WT cells controls. Triple KO cells were identified sequentially by selecting CD162^−^ cells, followed by CD43^−^ cells, and finally CD44^−^ cells. CLA expression was then assessed within the CD162^−^CD43^−^CD44^−^ population. For control samples, each marker was analyzed relative to the total live-cell population rather than sequential parent gates. **C)** Flow cytometry analysis of CLA depletion in T cells following triple CD162/CD43/CD44 KO compared with WT.

**Supplementary figure S6.**
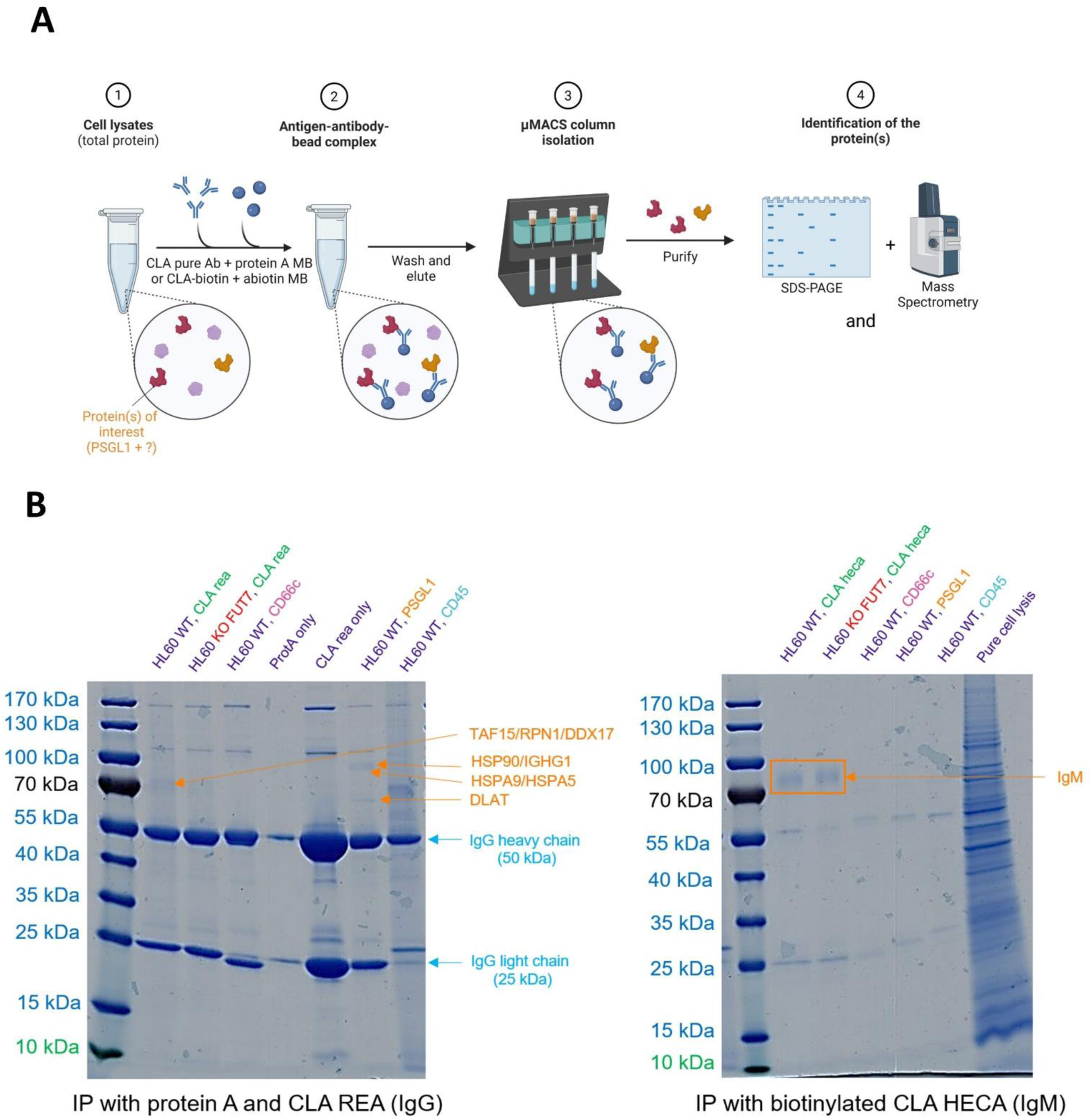
Identification of CLA antibody carrier proteins in HL60 cells. **A)** Workflow of the immunoprecipitation (IP) protocol. Two IP approaches were performed using HL60 cell lysates: (i) purified CLA REA antibody (IgG) coupled to Protein A, or (ii) biotinylated CLA HECA-452 antibody (IgM) coupled to anti-biotin microbeads. Immunoprecipitated proteins were analyzed by SDS-PAGE and identified by peptide mass fingerprint (PMF) analysis. Created with BioRender.com. **B)** SDS-PAGE analysis of IP samples from HL60 WT or CLA-depleted FUT7 KO cells. Left: IP performed with purified antibodies and Protein A. Right: IP performed with biotinylated antibodies and anti-biotin microbeads. Controls included CD66c antibody, Protein A only, CLA REA only, PSGL1/CD162 antibody, and CD45 antibody. Besides IgG-associated bands, the objective was to identify in HL60 WT cells treated with CLA antibodies: (i) a CD162/PSGL1 band comparable to the PSGL1/CD162 control, and (ii) additional bands potentially corresponding to alternative CLA carrier proteins that were absent in FUT7 KO cells. However, no clear candidate bands were detected. The only visible but weak band following CLA REA IP was identified by PMF as TAF15, RPN1 and DDX17 (TATA-box binding protein-associated factor 15, ribophorin-1 and DEAD-box helicase 17), likely representing nonspecific binding artifacts rather than CLA carrier proteins. A ~60 kDa band observed in the PSGL1/CD162 condition was identified as DLAT (dihydrolipoamide S-acetyltransferase) by PMF analysis. Additional weak bands corresponded to heat shock proteins (HSPs) and IGHG1 (immunoglobulin heavy constant gamma 1). Following IP with biotinylated antibodies, no specific bands were detected apart from bands corresponding to the IgM antibody identified by PMF analysis.

**Supplementary figure S7.**
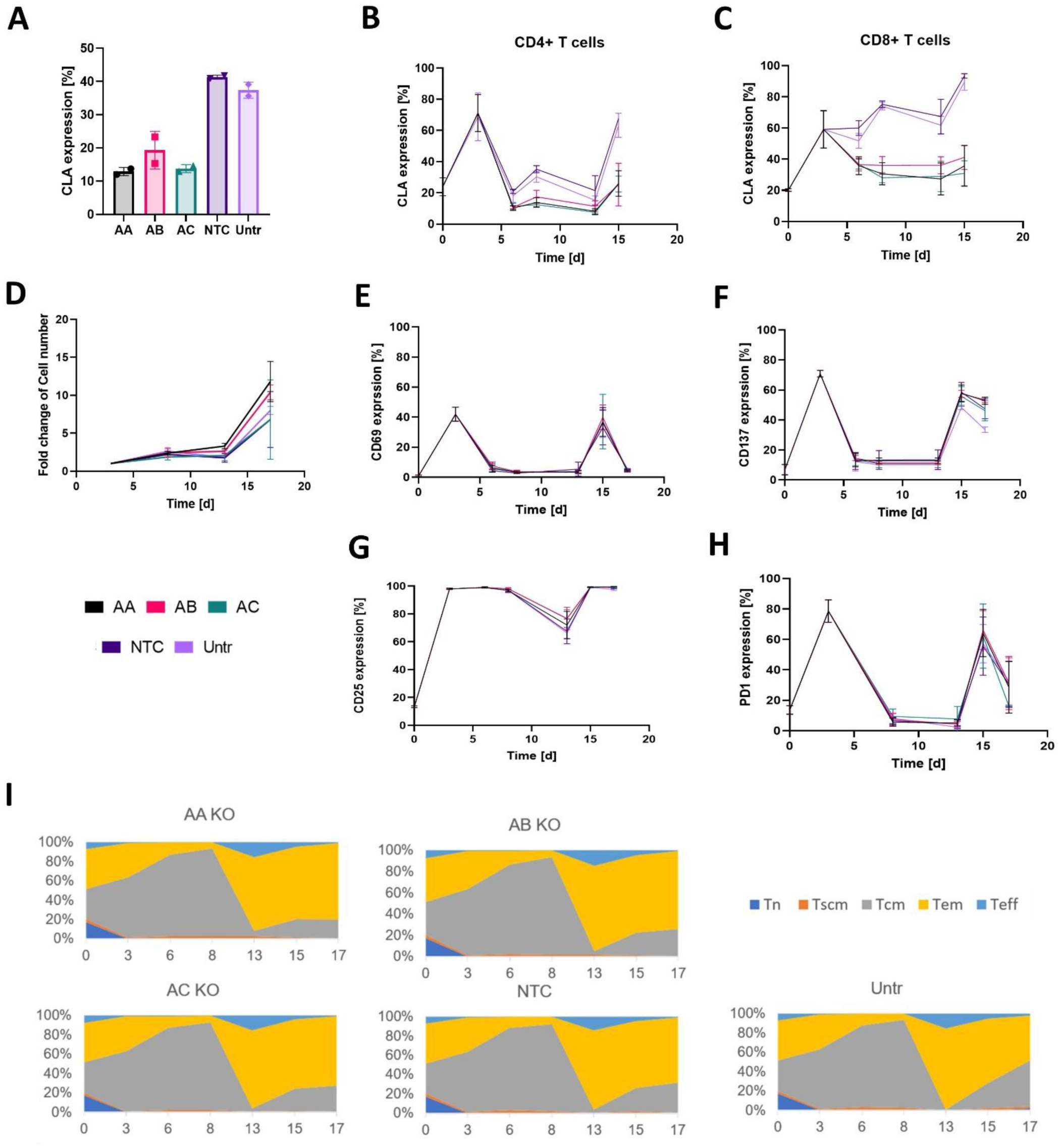
Analysis of FUT7 KO influence on T cell proliferation, markers and phenotype. **A)** Bar plot showing CLA expression on peripheral T cells on day 8 after FUT7 KO using three sgRNAs (AA, AB, AC) compared with non-transduced electroporated control (NTC) and untreated control (Untr). **B-C)** CLA expression over time on CD4^+^ **B)** and CD8^+^ **C)** T cells after FUT7 KO. **D)** Fold change in peripheral T cell counts over time following FUT7 KO. **E-H)** Expression of activation and exhaustion markers on T cells over time: CD69 **E)**, CD137 **F)**, CD25 **G)**, and PD-1 **H)** post-FUT7 KO. **I)** Area plots showing T cell differentiation after FUT7 KO. Percentages of T cell phenotypes over time were determined by flow cytometry based on CD62L, CD45RA and CD95 expression to assess differentiation status. **A-I)** Cells were activated with TransAct™ on days 0 and 13. All assays were performed in duplicate using blood from two independent donors. Tn: naïve T cells. Tscm: memory stem cell T cells. Tcm: central memory T cells. Tem: effector memory T cells. Teff: effector T cells.

**Supplementary figure S8.**
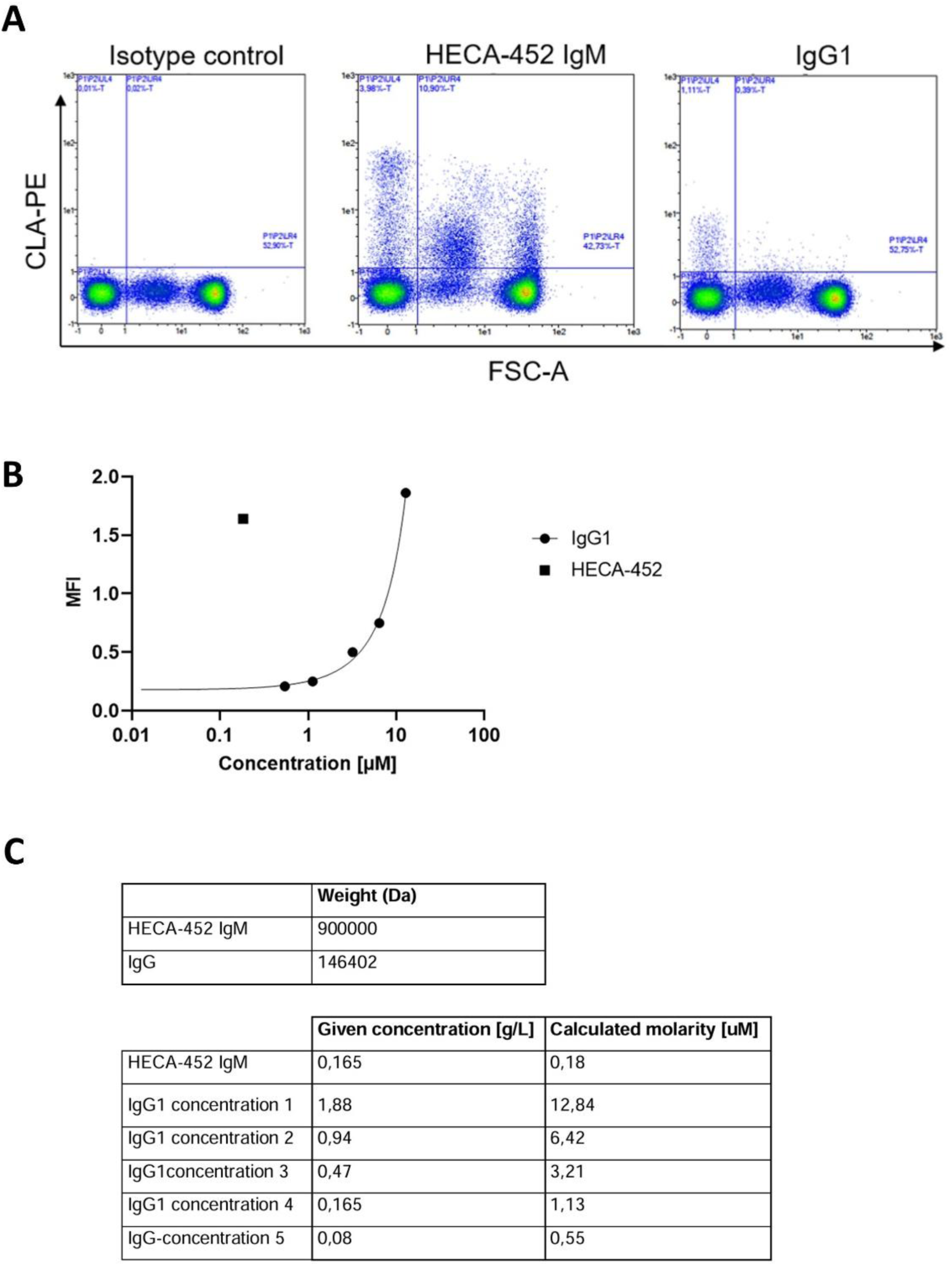
Titration of CLA REA IgG and HECA-452 IgM antibodies to compare staining performance. **A)** Density plot of flow cytometry showing binding of 0.165 g/L CLA REA IgG or HECA-452 IgM antibodies on PBMCs. **B)** Graph of mean fluorescence intensity (MFI) across a range of antibody concentrations. **C)** Calculation of the molarity of the anti-CLA HECA-452 IgM and the IgG. The weight of the HECA-452 IgM is estimated based on similar antibodies. Both antibodies were evaluated for their staining performance toward CLA at the indicated concentrations. The respective molarity was calculated with the estimated weight.

**Supplementary figure S9.**
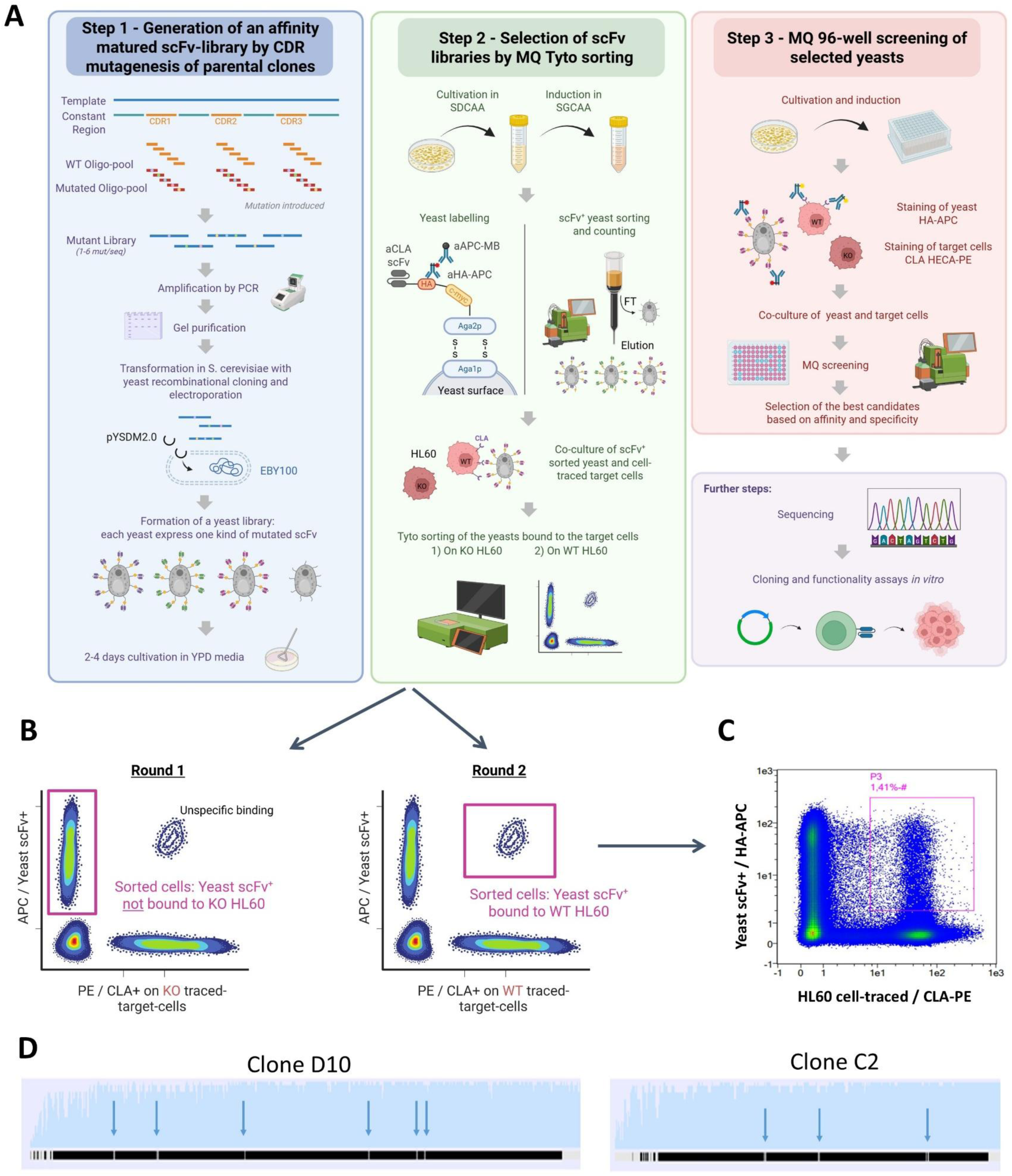
Affinity maturation by yeast surface display (YSD). **A)** Workflow of the in vitro YSD experiment. **B)** Schematic of the selection strategy during Tyto sorting. Round 1: HL60 KO cells are used to remove yeast expressing the HA tag that binds non-specifically. Round 2: HL60 WT cells are used to select yeast expressing the HA tag that specifically bind target cells. **A-B)** Created with BioRender.com. **C)** Flow cytometry analysis of yeast after Round 2 sorting. 1.41% of cells were selected, consistent with typical yields for this procedure. **D)** Sequencing results of two positive clones, D10 and C2, containing 6 and 3 mutations indicated with blue arrows, respectively. Clone C2 had a truncated heavy chain, so predicted mutations were only identified in the light chain.

**Supplementary figure S10.**
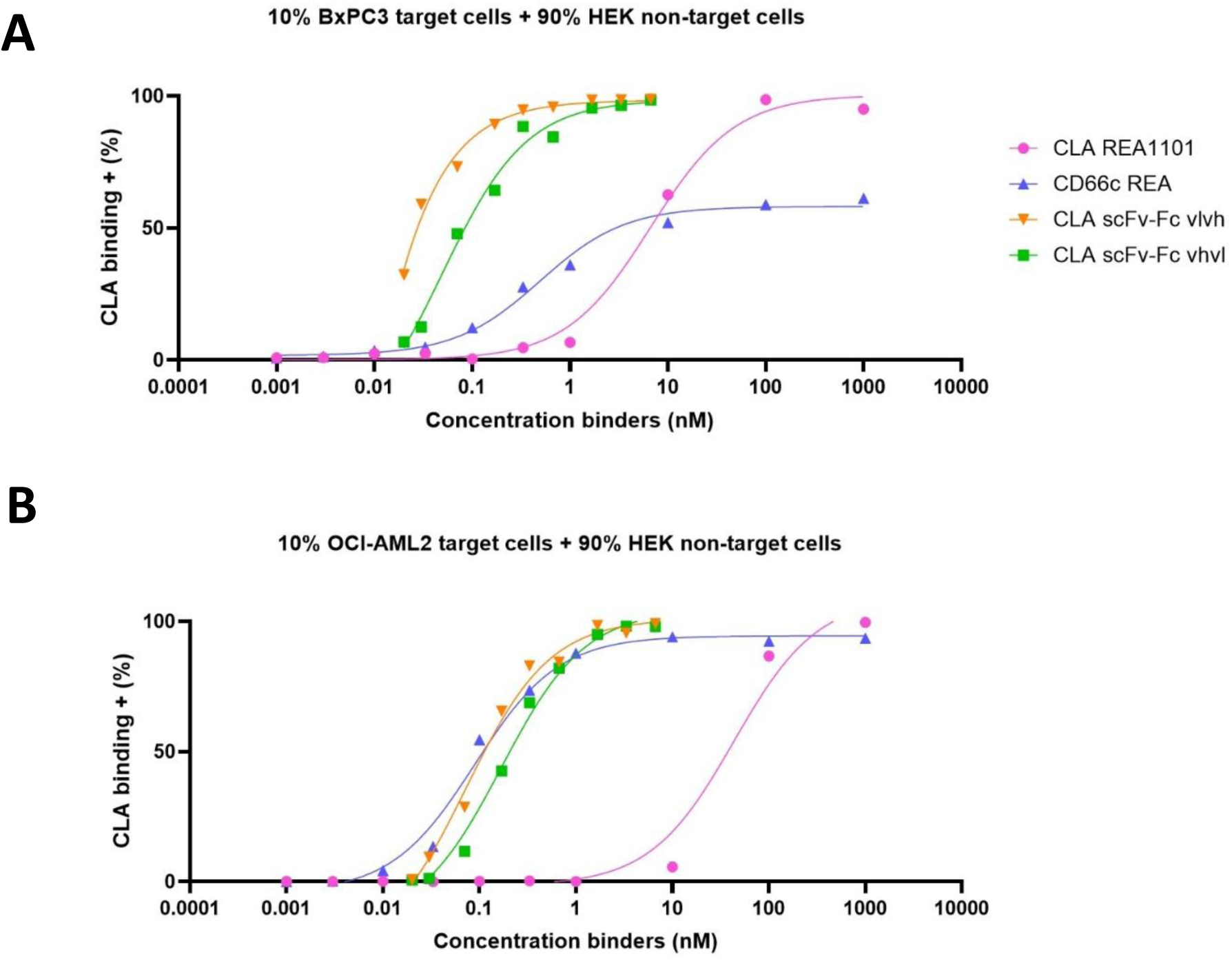
Binding performance of CLA scFv-Fc (VHVL or VLVH) compared with CLA REA IgG and CD66c REA IgG controls, evaluated by flow cytometry. 10% of target cells (A: BxPC3, B: OCI-AML2) were mixed with 90% non-target HEK cells. Data was analyzed using a three-parameter inhibitor-versus-response model with simple logistic regression in GraphPad Prism.

**Supplementary figure S11.**
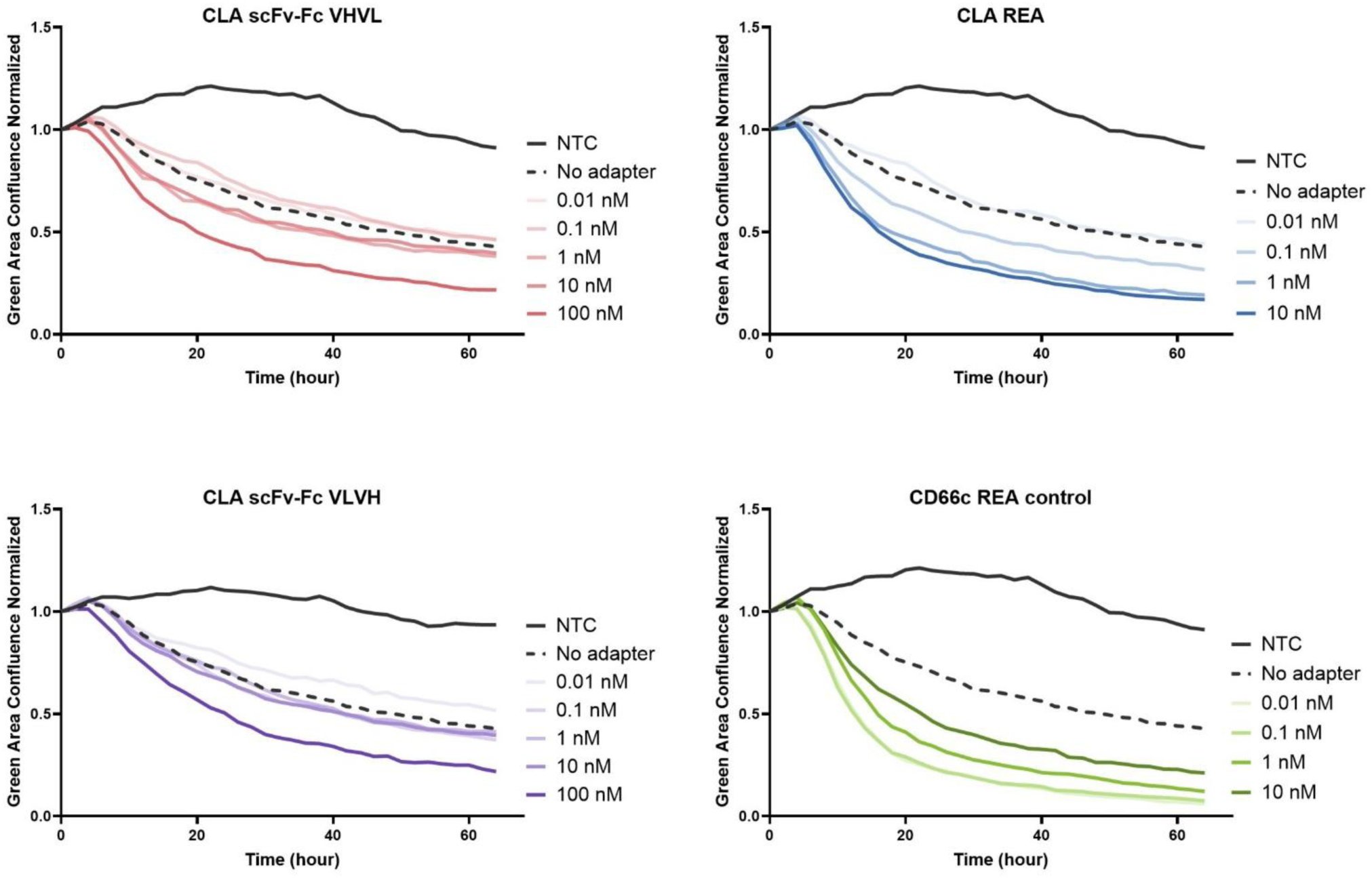
Killing curves of CLA scFv-Fc AdCAR assay for a second independent donor. GFP-based BxPC3 tumor cell confluence, normalized to the first time point, is shown for CLA AdCAR constructs at an E:T ratio of 2:1 using: CLA scFv-Fc VHVL adapter, CLA scFv-Fc VLVH adapter, CLA REA adapter or CD66c REA adapter. Results are compared with non-transduced T cells (NTC) and AdCAR without adapter (no-adapter) controls. Biotinylated adapters were tested at 0.01-100 nM for scFv-Fc adapters and 0.01-10 nM for REA controls.

**Supplementary figure S12.**
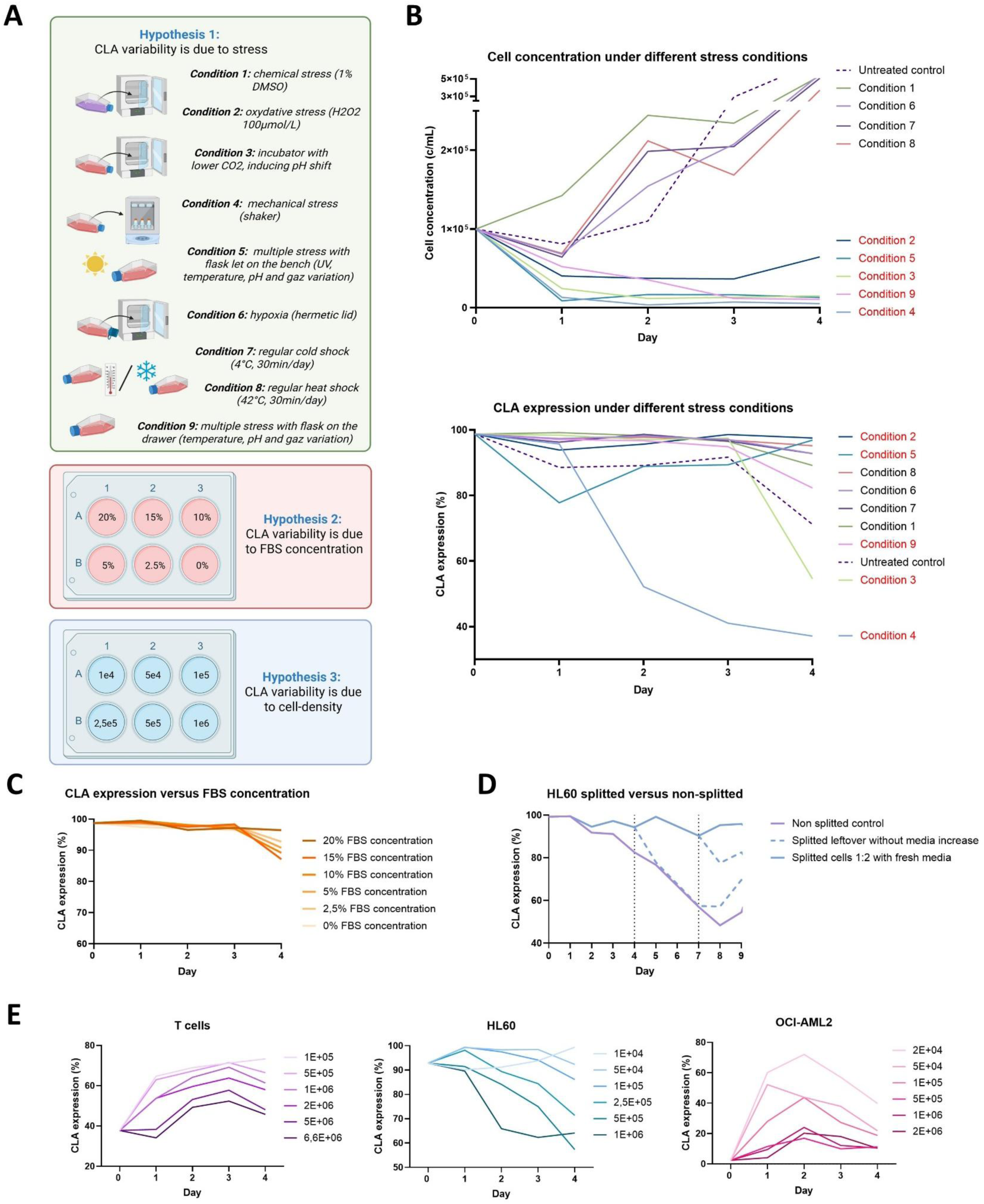
Investigation of CLA variability under three hypotheses: stress response, FBS concentration and cell density. **A)** Schematic of experimental conditions testing the impact of stress, FBS or cell density on CLA expression in HL60 cells. Created with BioRender.com. **B)** CLA expression and cell concentration over time in HL60 cells under various stress conditions. Conditions marked in red indicate treatments that induced widespread apoptosis or cell death. **C)** CLA expression over time in HL60 cells cultured with varying FBS concentrations (0-20%). **D)** CLA expression over time in HL60 cells seeded at 2.5 × 10^5^ cells/mL under different splitting conditions on days 4 and 7: non-split control, 1:2 split with fresh media, and split leftovers without fresh media. Split cells showed reduced CLA expression compared to non-split controls. **E)** CLA expression over time at different starting cell densities in T cells, HL60 and OCI-AML2 cell lines.

**Supplementary figure S13.**
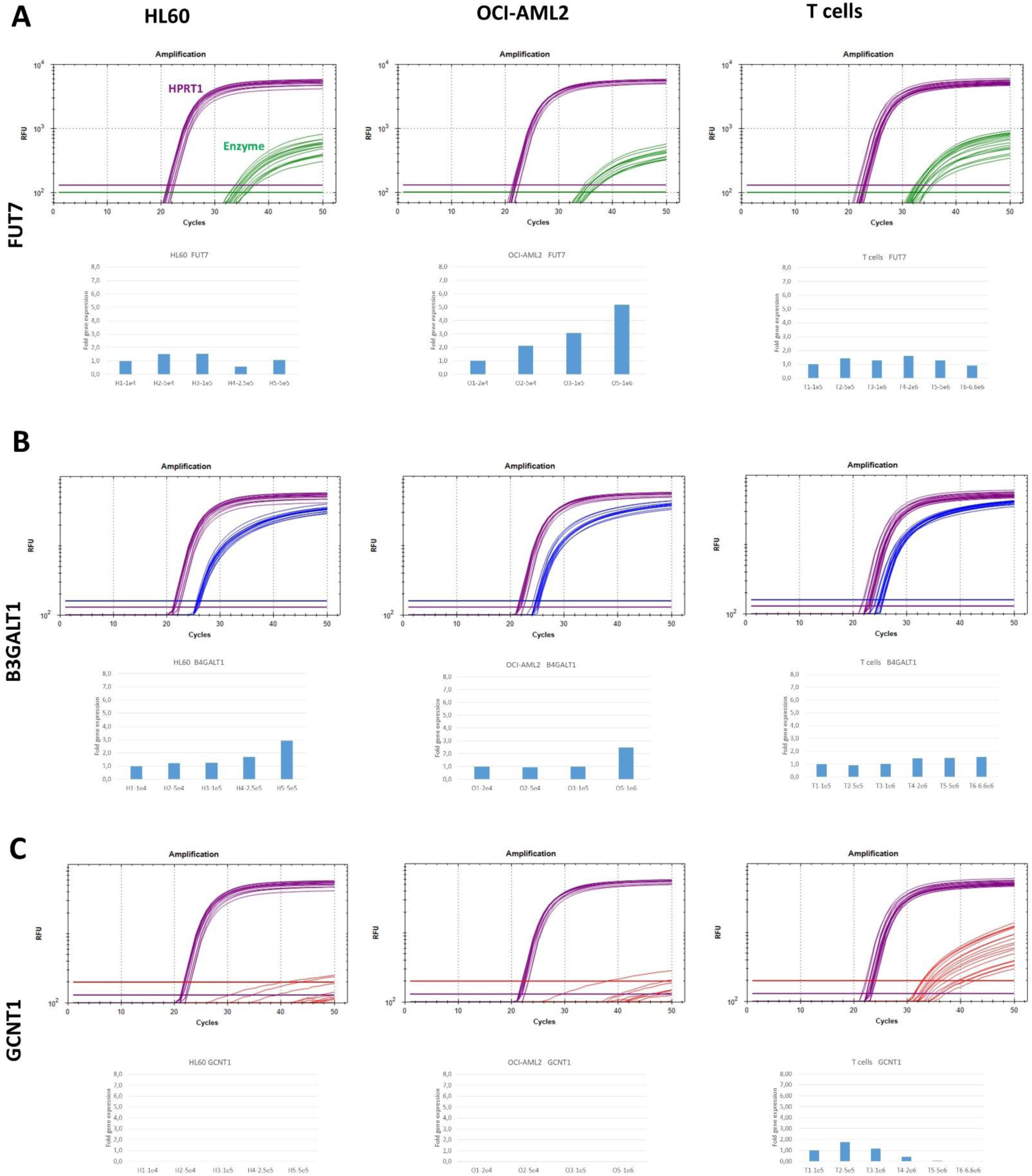
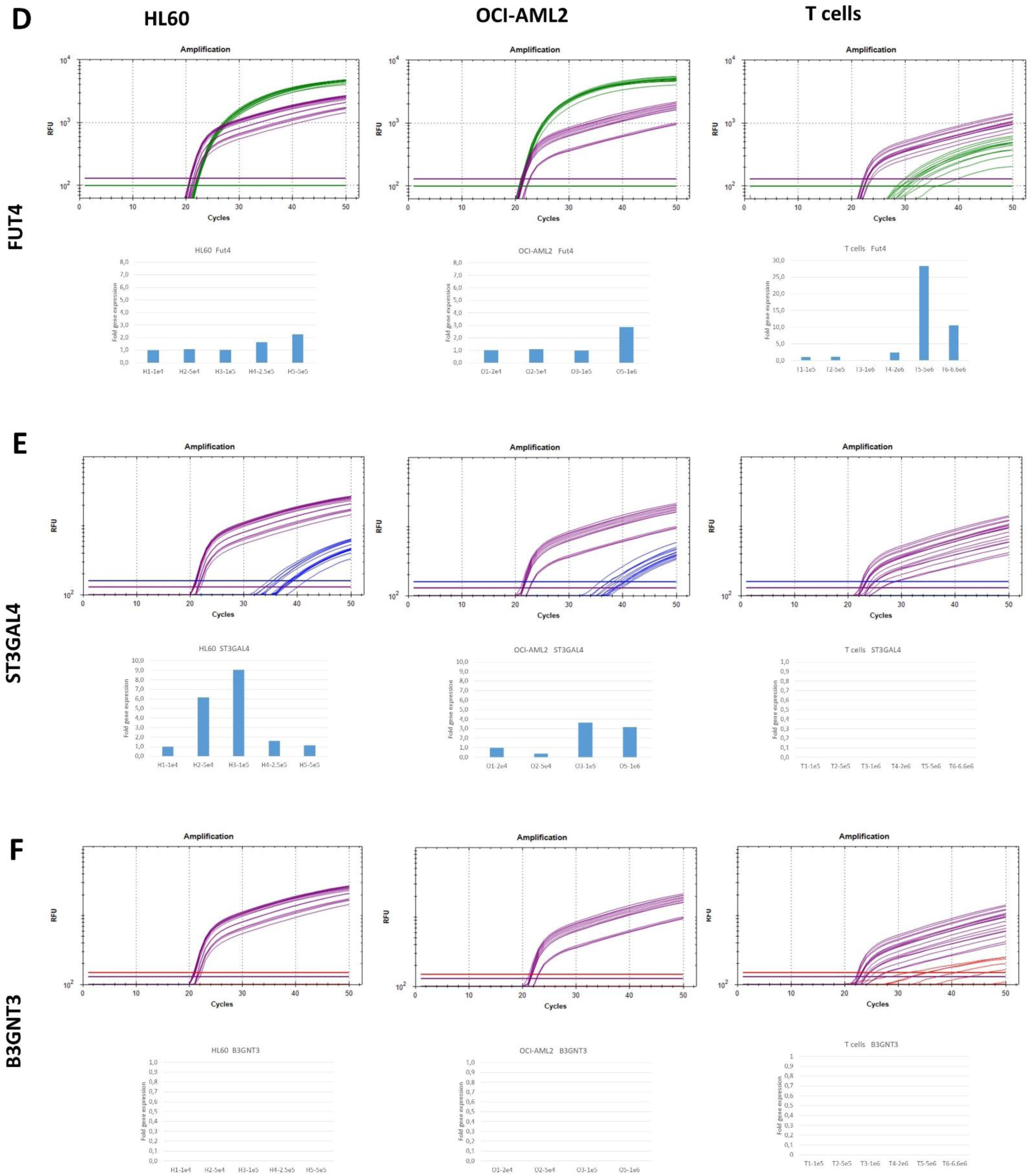
RT-qPCR analysis of enzymes involved in CLA biosynthesis in multiple cell lines after 5 days of culture at different cell densities. Cell lines analyzed: HL60, OCI-AML2 and peripheral T cells. **A-F)** Expression of key enzymes in CLA biosynthesis: **A)** α1,3-fucosyltransferase 7 (FUT7), **B)** β1,3-galactosyltransferase 1 (B3GALT1), **C)** β1,6-N-acetylglucos-aminyltransferase 1 (GCNT1), **D)** α1,3-fucosyltransferase 4 (FUT4), **E)** β-galactoside α2,3-sialyl-transferase 4 (ST3GAL4), **F)** β1,3-N-acetylglucos-aminyltransferase 3 (B3GNT3). For each enzyme and cell line, the amplification profile (cycle number versus relative fluorescence units [RFU]) is shown to illustrate PCR efficiency, with the HPRT1 housekeeping gene in purple. The bar plots below show the fold change in enzyme expression at the different cell densities. Fold changes were calculated using the delta-delta Ct method and normalized to HPRT1 housekeeping gene.

**Supplementary figure S14.**
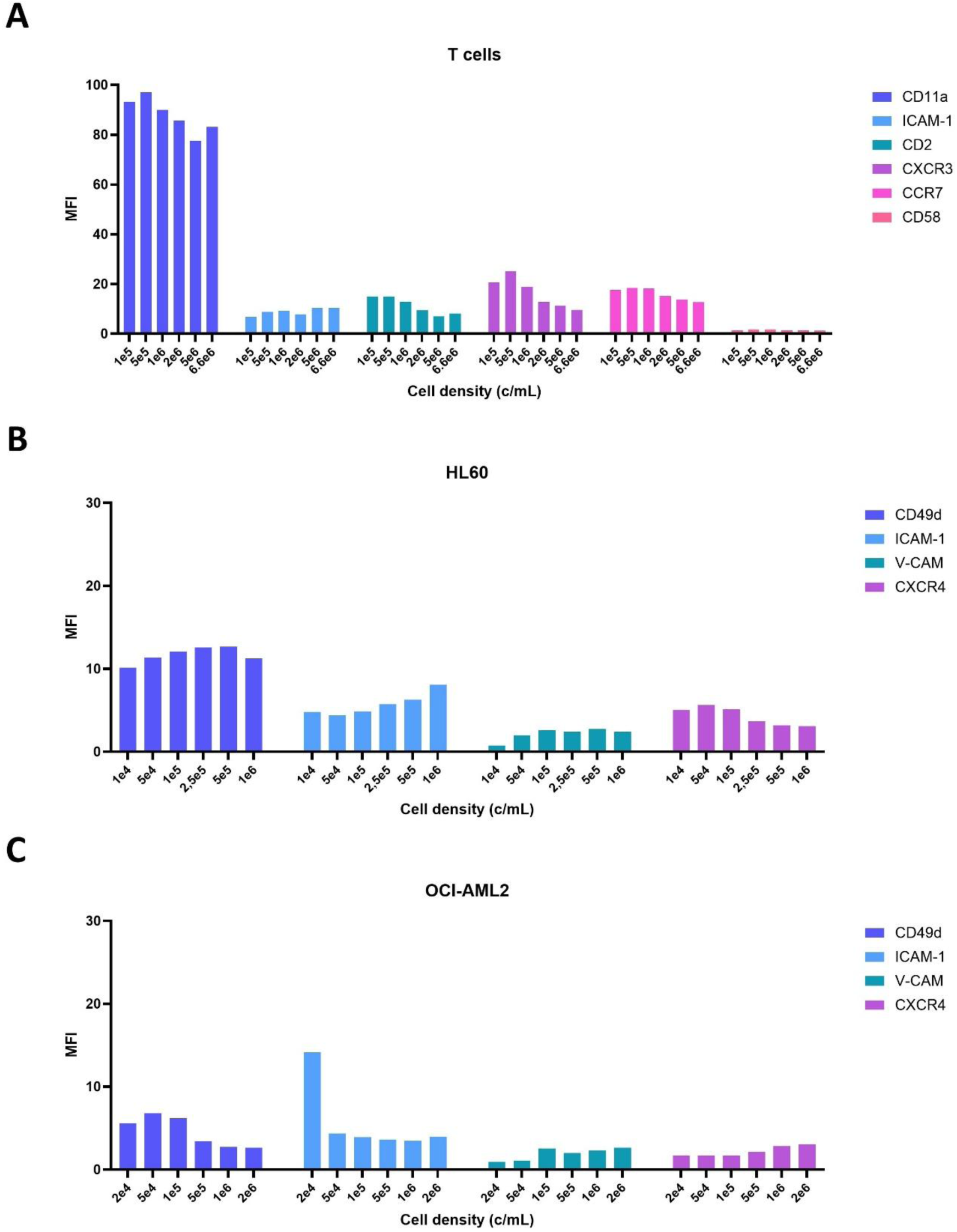
Flow cytometry analysis of adhesion molecule regulation at varying cell densities. Expression of adhesion molecules was assessed by mean fluorescence intensity (MFI) on T cells **A)**, HL60 cells **B)** or OCI-AML2 cells **C).** Cells were cultured for four days at densities ranging from 1 × 10^4^ to 6.6 × 10^6^ cells/mL, depending on the cell type, and analyzed by flow cytometry.

**Supplementary figure S15.**
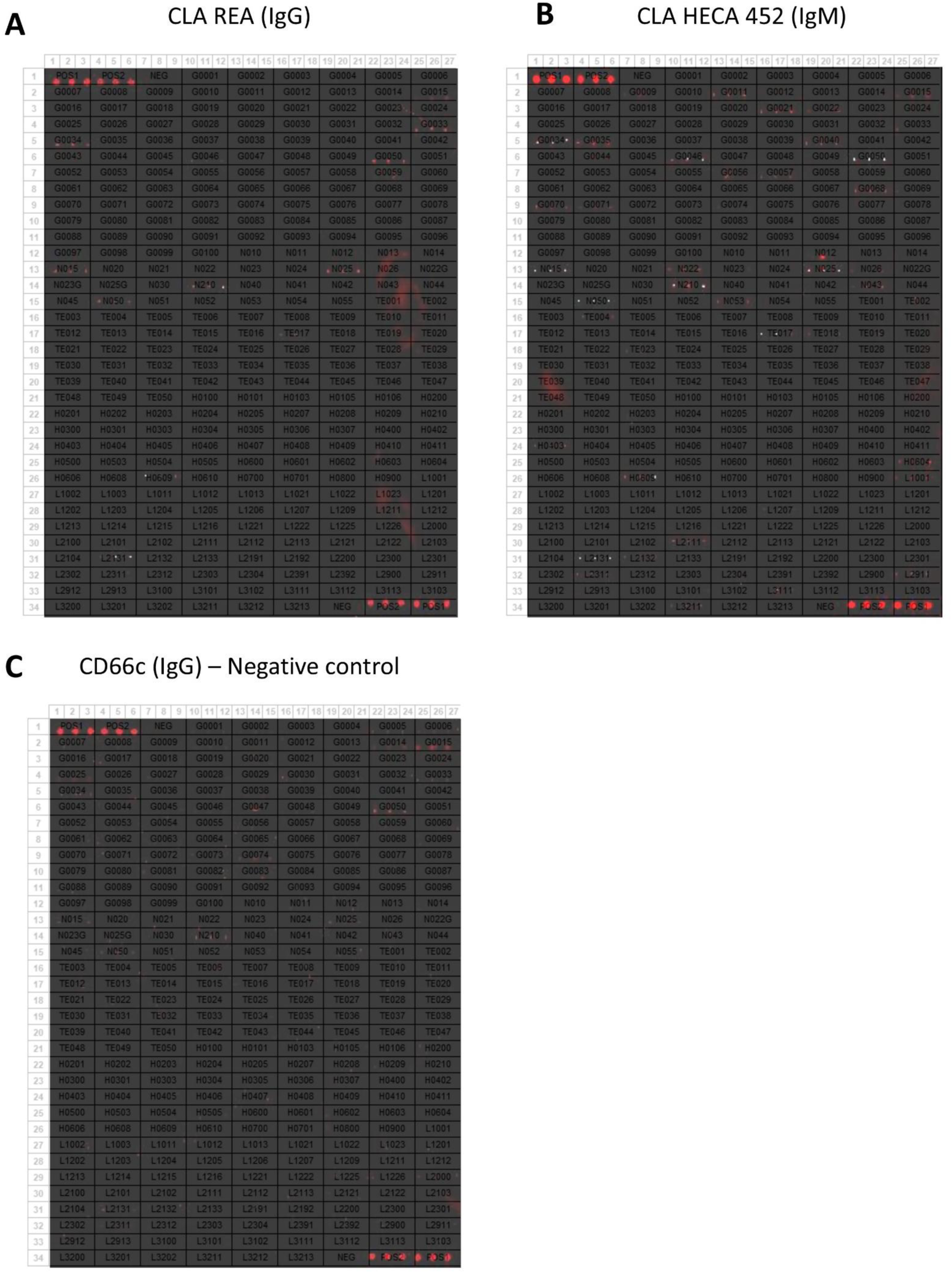
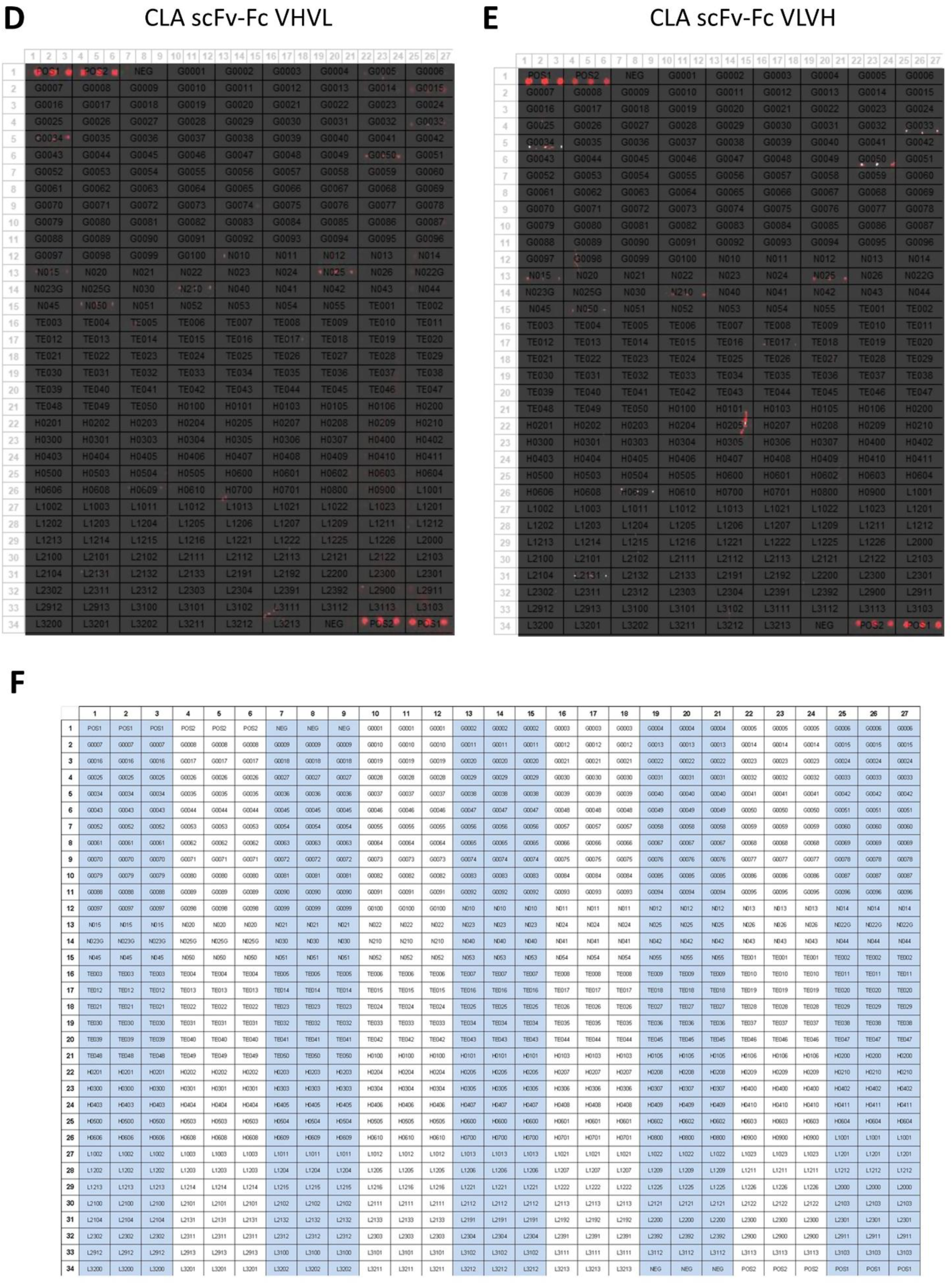
300-glycan array scan images showing luminescent binding signals overlaid with the plate layout. The assay was performed using **A)** CLA REA, **B)** CLA HECA-452, **C)** CD66c (background control), **D)** CLA scFv-Fc VHVL, and **E)** CLA scFv-Fc VLVH. **F)** Corresponding plate layout. Each sample was tested in triplicates. Raw and processed data, as well as full glycan names, are provided in the supplementary materials.

